# A chromosome-level assembly of the “Cascade” hop (Humulus lupulus) genome uncovers signatures of molecular evolution and improves time of divergence estimates for the Cannabaceae family

**DOI:** 10.1101/2022.03.24.485698

**Authors:** Lillian K Padgitt-Cobb, Nicholi J Pitra, Paul D Matthews, John A Henning, David A Hendrix

**Affiliations:** Department of Biochemistry and Biophysics, Oregon State University, Corvallis, Oregon; Department of Research and Development, Hopsteiner, S.S. Steiner, Inc., 1 West Washington Avenue, Yakima, Washington 98903, United States; Forage Seed and Cereal Research Unit, USDA-ARS, Corvallis, Oregon; Department of Crop and Soil Science, Oregon State University, Corvallis, Oregon; School of Electrical Engineering and Computer Science, Oregon State University, Corvallis, Oregon

**Keywords:** Cannabaceae, *Humulus lupulus*, Genome assembly, Molecular evolution, Whole genome duplication, gene families, synteny

## Abstract

- We present a chromosome-level assembly of the Cascade hop (Humulus lupulus L. var. lupulus) genome. The hop genome is large (2.8 Gb) and complex, and early attempts at assembly resulted in fragmented assemblies. Recent advances have made assembly of the hop genome more tractable, transforming the extent of investigation that can occur.
- The chromosome-level assembly of Cascade was developed by scaffolding the previously-reported Cascade assembly generated with PacBio long-read sequencing, and polishing with Illumina short-read DNA sequencing. We developed gene models and repeat annotations, and used a controlled bi-parental mapping population to identify significant sex-associated markers. We assess molecular evolution in gene sequences, gene family expansion and contraction, and time divergence using Bayesian inference.
- We identified the putative sex chromosome in the female genome based on significant sex-associated markers from the bi-parental mapping population. While the estimate of repeat content (~64%) is similar to the hemp genome, syntenic blocks in hop contain a greater percentage of LTRs. Hop is enriched for disease resistance-associated genes in syntenic gene blocks and expanded gene families.
- The Cascade chromosome-level assembly will inform cultivation strategies and serve to deepen our understanding of the hop genomic landscape, benefiting hop researchers and the Cannabaceae genomics community.

## Introduction

Hop (*Humulus lupulus* L. var. lupulus) is a diploid (2n=18+XX/XY), wind-pollinated perennial plant (Small, 1997; Smith *et al.*, 2006) with cultural, economic, and pharmacological significance, including use in brewing and consumables for flavor and aroma. *Humulus lupulus* is typically dioecious, having both male and female plants, although monoecious individuals also occur (Edwardson, 1952). Hop grows optimally between the 35° and 55° latitude in the Northern and Southern hemispheres (Barth *et al.*, 1994; McCoy *et al.*, 2019); however, countries at lower latitudes also produce hops (McCoy *et al.*, 2019). The female inflorescences, or cones, of hop plants are known as “hops,” and contain lupulin glands (glandular trichomes), which are the primary site of synthesis and storage of resins, bitter acids, essential oils, and flavonoids (Zanoli & Zavatti, 2008; Wang *et al.*, 2008; Korpelainen & Pietiläinen, 2021) (Figure 1a).

**Figure 1.**
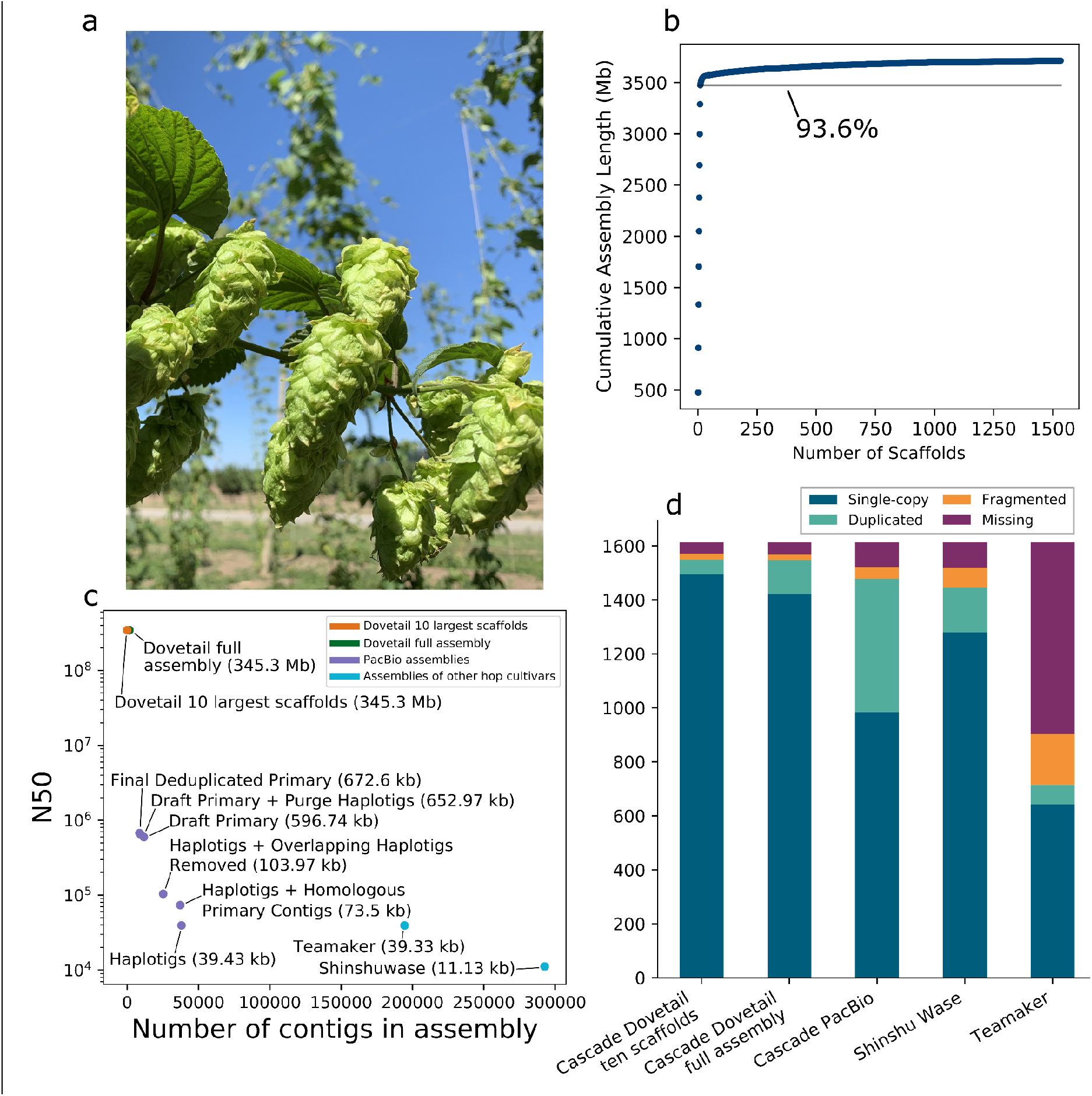
Analysis of Cascade Hop Dovetail assembly. **a)** Image of hop cones. **b)** Scatter plot depicting the cumulative assembly length on the y-axis relative to the number of scaffolds on the x-axis, showing that 93.6% of the assembly is contained in the largest ten scaffolds. **c)** Scatter plot showing the number of contigs or scaffolds in a given assembly on the x-axis versus the N50 on the y-axis. **d)** BUSCO result comparison among the assemblies for all Dovetail scaffolds; the ten largest scaffolds in the Dovetail assembly; PacBio Cascade; Shinshu Wase; and Teamaker.

Hop cultivar Cascade is known for its floral and citrus aroma, and is the most widely produced American “aroma” hop (Hieronymus, 2012). Cascade was developed at Oregon State University and USDA (USDA-ARS; Horner *et al.*, 1972). The pedigree of Cascade is [Fuggle × (Serebrianka × Fuggle-seedling)] × OP (open-pollinated seed) (Henning *et al.*, 2004). The oil content of Cascade is rich in myrcene (45-60% of oil content), cohumulone (33-40% of oil content), and humulene (8-13% of oil content) (Hieronymus, 2012). Linalool and geraniol also contribute to the flavor and aroma of Cascade (Takoi *et al.*, 2009).

### Evolutionary history of *Humulus lupulus*

*Humulus* originated in East Asia in a region corresponding to modern-day China (Neve, 1976; Korpelainen & Pietiläinen, 2021) and dispersed from East Asia into Central-West Eurasia approximately 25.4-10.7 mya (Jin *et al.*, 2020). The emergence of eastern and western lineages of *Humulus* was a result of a geographical barrier that occurred with the rise of the Tibetan Plateau and the emergence of the dry climate of northern Central Asia (Jin *et al.*, 2020). *Humulus* migrated westward to Europe via the Caucasus mountains (Murakami *et al.*, 2006), and migrated from East Asia to North America via the land bridge, Beringia (Liu *et al.*, 2002). After crossing Beringia, *Humulus* diverged into three North American varieties: *H. lupulus* var. *neomexicanus* Nelson and Cockerell, *H. lupulus* var. *pubescens* E. Small, and *H. lupulus* var. *lupuloides* E. Small (Murakami *et al.*, 2006; Tembrock *et al.*, 2016; McCallum *et al.*, 2019; Korpelainen & Pietiläinen, 2021).

*H. lupulus* is separated into five taxonomic varieties based on morphology and geography that can be hybridized to produce fertile offspring (McCoy *et al.*, 2019): var. *lupulus* L., var. *cordifolius* (Miguel) Maximowicz, var. *neomexicanus*, var. *pubescens*, and var. *lupuloides*. *H. lupulus* var. *lupulus* L. is native to Europe and Asia, and is the primary source of germplasm for the majority of hop cultivars, and is also naturalized in many regions of the world as a result of escape from commercial hop yards. *H. lupulus* var. *cordifolius* is a variety of wild hops found in Japan; *H. lupulus* var. *neomexicanus* is endemic to the North American Cordillera; var. *pubescens* grows in the midwestern United States and Canadian provinces; and *H. lupulus* var. *lupuloides* grows in central and eastern North America (Small, 1978, 1981; Reeves & Richards, 2011; McCoy *et al.*, 2019). The geographical distribution of *Humulus* varieties is also consistent with the flavonoid dichotomy, defined as the presence or absence of 4′-*O*-methylchalcones (Stevens *et al.*, 2000; Hummer *et al.*, 2004).

Wild hops are not used in brewing due to bitterness and unpleasant aroma; however, wild varieties provide a repository of genetic diversity to develop new cultivars with enhanced resistance to abiotic and biotic stresses. Well-known crosses between North American and European hop varieties include Brewer’s Gold and Comet (Tembrock *et al.*, 2016). Further, North American wild hops have high α-acid content and resistance to *Verticillium* wilt disease (Neve, 1991).

Within Cannabaceae, *Humulus* and *Cannabis* form a sister clade to the clade containing *Celtis*, *Trema*, and *Parasponia. Parasponia* is nested within the non-monophyletic *Trema* (Yang *et al.*, 2013; Sun *et al.*, 2016; Jin *et al.*, 2020). *Humulus lupulus* is one of three species of *Humulus*, along with *H. yunnanensis* Hu. and *H. japonicus* Siebold & Zucc. (synonymous with *Humulus scandens* (Lour.) Merr. (‘Humulus scandens (lour.) Merr’, 2019). We will refer to *Humulus scandens* herein as *Humulus japonicus*.

Phylogenetic analysis of modern and ancient *Cannabis* samples revealed that *H. japonicus* is more closely related to both ancient and modern samples of *Cannabis*, than *H. lupulus* (Грудзинская, 1988; Mukherjee *et al.*, 2008). Successful grafting between *C. sativa*, *H. japonicus*, and *H. lupulus* further underscores the close relationship between these species (Crombie & L. Crombie, 1975). Further, the xanthohumol pathway in hop and the cannabinoid pathway in *Cannabis* contain enzymes that perform analogous reactions (Page & Nagel, 2006).

The date of divergence of *Humulus* and *Cannabis* has been a source of debate (He *et al.*, 2013; McPartland, 2018; Jin *et al.*, 2020). Unraveling the complex evolutionary history of species in the Cannabaceae has been challenged by sparse fossil evidence and genomic resources. With the development of new genomic data, previously estimated divergence dates can be reevaluated and refined (Wilkinson *et al.*, 2011; Herendeen *et al.*, 2017; Silvestro *et al.*, 2021).

### Diseases impacting hop production

Hop is susceptible to fungal diseases, including powdery mildew (*Podosphaera macularis*), downy mildew (*Pseudoperonospora humuli* (Miyabe & Takah.) G.W. Wilson), and black root rot (*Phytophthora citricola* Sawada). Hop is also impacted by many viruses, including *Hop latent virus* (HpLV), *American hop latent virus* (AHLV), *Hop mosaic virus* (HpMV), and *Apple mosaic virus* (ApMV) (McCoy *et al.*, 2019).

### Motivation for a chromosome-level genome assembly

The hop genome is large and heterozygous. The size and complexity of the hop genome challenged previous assembly efforts (Natsume *et al.*, 2015; Hill *et al.*, 2017; Padgitt-Cobb *et al.*, 2021). Recent advances, including the development of long-read sequencing and haplotype-aware assembly algorithms, have made assembly of the hop genome tractable, and provide a transformational foundation for plant breeding and genomics. Development of cultivars with enhanced tolerance to abiotic and biotic stresses, short-stature, and distinct flavor and aroma profiles is a research area of priority (McCoy *et al.*, 2019) and would be benefitted by further development of genomic resources.

Here we describe a chromosome-level assembly of the Cascade hop genome developed with the draft, phased PacBio long-read assembly of Cascade (Padgitt-Cobb *et al.*, 2021) and Dovetail high-throughput chromatin conformation capture (Hi-C) libraries. Hi-C allows detection of long-range DNA interactions by sequencing fragments of cross-linked chromatin, providing information about spatial organization of chromatin (Dudchenko *et al.*, 2017). Using this long-range information, the PacBio contigs can be ordered and oriented into chromosome-level scaffolds.

## Materials and Methods

### Genome sequencing and assembly

Hi-C libraries were used to scaffold and correct the contigs from the PacBio assembly long-read assembly (Padgitt-Cobb *et al.*, 2021) using HiRise (Putnam *et al.*, 2016) by Dovetail Genomics. DNA was extracted from Cascade leaves using a method previously described (Padgitt-Cobb *et al.*, 2021), which involved a modified CTAB method to reduce the inclusion of small, sheared DNA fragments. Dovetail scaffolds were polished with a set of 563,456,691 DNA short reads from Cascade using the Polca polishing tool (Zimin & Salzberg, 2020). We used samtools-1.11 flagstat to assess the mapping rate of the DNA short reads to the Dovetail assembly. We used BUSCO v5.2.2 (Waterhouse *et al.*, 2018) to assess the assembly completeness, which also incorporated Augustus-3.3.2 (Stanke *et al.*, 2006) and both Embryophyta and Viridiplantae databases from OrthoDB v10 (Kriventseva *et al.*, 2019).

### Repeat annotation

We identified repeat sequences using an approach described previously (Padgitt-Cobb *et al.*, 2021). Briefly, we developed a *de novo* set of long terminal retrotransposon (LTR) annotations using suffixerator (GenomeTools) 1.6.1 (Gremme *et al.*, 2013), LTRharvest (GenomeTools) 1.6.1 (Ellinghaus *et al.*, 2008), LTR_FINDER_parallel v1.1 (Ou & Jiang, 2019), and LTR_retriever v2.7 (Ou & Jiang, 2018). We used suffixerator to index the assembly, then LTRharvest and LTR_FINDER_parallel were used to identify LTRs, and finally, LTR_retriever was used to synthesize and refine the results of LTRharvest and LTR_FINDER_parallel. We identified non-LTR types of repeats from plants using a database from MIPS PlantsDB (Nussbaumer *et al.*, 2013). The *de novo* set of LTRs and the database of plant repeats were aligned to the assembly using RepeatMasker version 4.1.0 (Smit *et al.*, 2015). The repeat annotation pipeline was performed on each scaffold separately. The detailed pipeline for repeat annotation can be found at RepeatAnnotationPipeline.md on GitHub.

### Gene model development

We developed our set of gene models with Transdecoder-v5.5.0 (Haas *et al.*, 2013) and MAKER (Holt & Yandell, 2011; Campbell *et al.*, 2014). In cases of overlapping gene models, we assigned preference to the Transdecoder gene models, because Transdecoder directly incorporates transcript expression evidence into gene model development. We incorporated RNA-seq data from lupulin glands and leaf (see Supplementary Information); leaf, meristem, and stem tissues (Padgitt-Cobb et al., 2021), as well as from hop cones during critical developmental stages (Eriksen et al., 2021). Putative gene functions were assigned based on similarity to known proteins from UniProt (accessed 08/24/2020 using search term: taxonomy:“Embryophyta [3193]” AND reviewed:yes) (Acids research & 2021, 2021) and Pfam protein domains (Pfam release 33.1) (Mistry *et al.*, 2021). The detailed pipeline for generating the gene models and then assigning putative function is provided in Supporting Information (SI) and on the GitHub project page in the directory ‘GeneModels’.

### Synonymous substitution rate (Ks)

To assess molecular evolution in hop and hemp, we calculated the synonymous substitution rate (Ks) (Li & Gojobori, 1983; Hughes & Nei, 1988; Hanada *et al.*, 2007) for anchor gene pairs identified with MCScanX (Wang *et al.*, 2012). Synonymous substitutions do not change the encoded amino acid (Kimura, 1977), and because substitution occurs at an approximately constant rate, the substitution rate can be treated as a proxy for elapsed time since the duplication of paralogous genes (Li, 1997; Vanneste *et al.*, 2015).

We visualized syntenic blocks on a genome-wide scale with SynVisio (Bandi & Gutwin, 2020), requiring a minimum MATCH_SIZE of 9, corresponding to at least ten anchor genes per block. We also ran MCScanX with default settings for downstream analysis, and visualized the blocks with Integrative Genomics Viewer (IGV) (Robinson *et al.*, 2011).

We performed a codon alignment for each anchor gene pair using MACSE alignSequences and exportAlignment (Ranwez *et al.*, 2018). We calculated Ks for each collinear gene pair individually (Xiao *et al.*, 2015) using the yn00 package within CodeML (Yang & Nielsen, 2000; Yang, 2007). We assessed functional enrichment in the syntenic blocks of hop vs hop and hop vs hemp with a hypergeometric test and performed a Benjamini-Hochberg multiple test correction (Benjamini & Hochberg, 1995) to obtain a q-value for each GO term (*FDR* < *0*.*05*).

### Estimation of divergence using Ks

Divergence dates are often estimated using Ks and the mutations per site per year (Koch *et al.*, 2000), denoted as λ, with the formula *T* = *Ks*/*2λ* (Koch *et al.*, 2000), where *T* is the length of time elapsed since the time of duplication or divergence (Lang *et al.*, 2018). A known divergence time is required to determine λ, which is reliant upon evidence from the fossil record (Wolfe *et al.*, 1987). There are different estimates of λ for plants, including 6.1 × 10^−9^ (Lynch & Conery, 2000) and 1.5 × 10^−8^ (Koch *et al.*, 2000) for *Arabidopsis*. In a previous study, the divergence date between *Humulus lupulus* and *Humulus japonicus* was estimated to be 3.74 mya based on Ks=*0*.*0157* ± *0*.*0056* using the *rbc*L sequence (Murakami, 2000), along with λ=2.1 × 10^−9^ (Wolfe *et al.*, 1987). In subsequent work, Murakami *et al.* used λ=1.23 × 10^−9^ (Xiang *et al.*, 2000) to calculate the divergence date for European, North American, and Asian hop lineages, and provided an updated estimate of the divergence date between *Humulus lupulus* and *Humulus japonicus* as 6.38 mya (Murakami *et al.*, 2006). We used λ=6.1 × 10^−9^ (Lynch & Conery, 2000; Fawcett *et al.*, 2009; He *et al.*, 2013), which was also used to calculate the divergence time between *Cannabis sativa* and *Morus notabilis* (He *et al.*, 2013).

We restricted Ks values to ≥ 0.01 and ≤ 2.0 (Vanneste *et al.*, 2015; Lang *et al.*, 2018), to limit the inclusion of Ks values from allelic variants (Kondrashov *et al.*, 2002; Parks *et al.*, 2018), and to avoid saturation at large Ks values (Barker *et al.*, 2008; Vanneste *et al.*, 2013; Lang *et al.*, 2018). Ks=0.01 corresponds to the approximate Ks value marking the divergence of *H. lupulus* and *H. japonicus* (Murakami, 2000).

We performed density estimation with log-transformed Ks values using Gaussian finite mixture modeling within the densityMclust function of mclust version 5.4.7 (Scrucca *et al.*, 2016) in R version 4.0.3 (Team & Others, 2013). Log-transformation allows better detection of peaks corresponding to duplication events (Lang *et al.*, 2018). We used the integrated complete-data likelihood (ICL) to select the number of clusters. ICL penalizes clusters that overlap excessively and tends to prefer clusters with well-delineated separation (Biernacki *et al.*, 2000). Our approach is adapted from previously described methods (Lang *et al.*, 2018; Li *et al.*, 2018; Shingate *et al.*, 2020; Mabry *et al.*, 2020; Ojeda-López *et al.*, 2020).

### Fossil-calibrated Time Tree

We used OrthoFinder version 2.5.2 (Emms & Kelly, 2019) to identify orthologous genes in eight species: *Cannabis sativa*, *Humulus lupulus*, *Morus notabilis*, *Parasponia andersonii*, *Prunus persica*, *Trema orientale*, *Vitis vinifera*, and *Ziziphus jujuba*. We collected single-copy orthologs present in all species and aligned the sequences with MACSE alignSequences and exportAlignment. Then, we extracted the third codon position from four-fold degenerate codon sites and concatenated the third positions to create a single alignment.

We used MCMCTree to perform a Bayesian estimate of divergence times, incorporating the concatenated alignment, the species-level phylogenetic tree from OrthoFinder, and fossil calibration data. We evaluated two molecular clock models with MCMCTree: strict molecular clock and independent log-normally distributed relaxed-clock model (clock=2) (Rannala & Yang, 2007), using a likelihood-ratio test (LRT) to compare molecular clock models (Yang *et al.*, 2000) with the equation *LRT* = −*2*(*ln*(*L*_*s*_) − *ln*(*L*_*g*_)). MCMCTree parameters are described in SI Methods.

### Gene family expansion and contraction

We investigated the expansion and contraction of gene family size in a phylogenetic context with CAFE (De Bie *et al.*, 2006). We incorporated our time tree, along with the size of each orthologous gene group from OrthoFinder. We required orthogroups to contain not more than 100 genes from a single taxon (Gold *et al.*, 2019). Trees were visualized with FigTree 1.4.4 (Rambaut, 2006-2018).

We assessed GO term functional enrichment in gene families that are expanded or contracted in hop. We applied a hypergeometric test to identify GO terms that are enriched among all genes and all species that occur in the expanded and contracted gene families. Then, we performed a Benjamini-Hochberg multiple test correction (Benjamini & Hochberg, 1995) to obtain a q-value for each GO term.

We collected the top functionally-enriched GO terms among all statistically significant GO terms (*FDR* < *0*.*05*). We visualized the top GO terms with *FDR* < *1e* − *5*, sorting by observed *k*. Depicted in Figure 5c and 5d are a collection of these top ten functionally-enriched Biological Processes GO terms. GO term associations are available for download at http://hopbase.cqls.oregonstate.edu/Downloads.php.

### Construction of the linkage map

A controlled bi-parental mapping population was developed by crossing the female hop line ‘Comet’ (Zimmermann *et al.*, 1975) by the male USDA germplasm accession 64035M. Seeds were vernalized, germinated, and grown in the greenhouse as previously reported (Henning *et al.*, 2015). The population used for this study consisted of 281 offspring and the corresponding parents. DNA extraction for all genotypes, library preparation for Illumina-based genotyping-by-sequencing (GBS), and Illumina sequencing were all performed as previously reported (Padgitt-Cobb *et al.*, 2019).

Genetic mapping was performed as previously described (Padgitt-Cobb *et al.*, 2019). Briefly, SNPtag data files were imported into Microsoft Excel and numerical values (0,1,2) were converted to A, H, B (representing AA, AB, BB). Segregation for each locus was obtained and tested for goodness-of-fit with predicted test-cross segregation (1:1) or F2-segregation (1:2:1) using chi-squared tests. Loci with significant chi-squared test statistics for goodness-of-fit for test-cross segregation (1:1) were combined with loci with significant test statistics for F2-segregation and subsequently combined into male and female data sets. These two data sets were imported as F2-formated files and separately analyzed in JoinMap 5.0 (VAN Ooijen, 2011). Genetic maps were estimated using both maximum likelihood (ML) and regression mapping (with two rounds of map estimation). Under ML mapping, loci with resulting high “NN Fit (cM)” (NN Fit values > 100) were excluded from map estimation, and map estimation was re-run until no further high NN Fit values were observed. The resulting male and female F2 genetic maps were subsequently joined into a single “CP-Method” dataset and genotypes were re-coded with CP-Method codes (i.e.: ll, lm, nn, np, hh, hk and kk). Regression mapping in JoinMap 5.0 using CP-Methods (again, with two rounds of map estimation) was subsequently performed to obtain the overall unbiased length of the genetic map. ML mapping using CP-Method was also performed to provide a more-accurate placement of markers for genetic map estimation. The resulting genetic distance between markers using ML was always overestimated. As a result, the ML map marker distances were adjusted downwards to match the overall genetic map estimate obtained via use of regression mapping in JoinMap 5.0.

## Results

### Genome sequencing and assembly

The size of the PacBio primary assembly used to anchor the scaffolds is 3,711,963,939 bp, and the resulting Dovetail Hi-C assembly is 3,712,781,139 bp (SI Table S1). The polished Dovetail assembly is 3,713,677,344 bp (SI Table S2). The library insert size distribution is shown in SI Figure S1, and the link density histogram for read position versus mate position is shown in SI Figure S2. The estimated physical coverage for Dovetail is 208.29X. The N50 increased from 673 kb for PacBio to 345.208 Mb for Dovetail (Figure 1c; Table 1). The scaffold N90 increased from 221 kb for PacBio to 185.170 Mb for Dovetail (SI Figure S3). The ten largest Dovetail scaffolds have a total length of 3.47 Gb, and 93.6% of the assembly is represented in the ten largest scaffolds (Figure 1b).

**Table 1.** Comparative assembly statistics

| Category | PacBio assembly | Dovetail Hi-C polished assembly (full assembly) |
| --- | --- | --- |
| Assembly size (bp) | 3,711,963,939 bp | 3,713,677,344 bp |
| Number of scaffolds | 8,661 | 1,533 |
| Number of scaffolds > 1kb | 8,661 | 1,533 |
| Largest scaffold (bp) | 8,249,941 bp | 476,374,791 bp |
| N50 (Mb) | 672,603 bp | 345,299,309 bp |
| N90 (Mb) | 221,394 bp | 185,200,997 bp |
| GC content (%) | 39.14% | 39.13% |
| Assembly gaps | 0 | 8172 |
| Percent of genome in gaps | 0 | 0.02% |

GC content of the full, polished Dovetail assembly is 39.13% and 0.022% Ns. Nucleotide content in the largest ten scaffolds is depicted in SI Figure S4a, while enrichment of genome-wide dinucleotide content is visualized as a heatmap in SI Figure S4b, and shows a depletion of CG content. Expected versus observed frequency of CHG and CHH trinucleotides, where H represents A, C, or T nucleotides, is shown in SI Figure S4c, revealing an enrichment of CHH trinucleotides. CHG and CHH trinucleotides are associated with DNA methylation, which is involved in gene regulation of essential plant processes, including growth and development (Zhang *et al.*, 2018b).

### Assembly completeness with BUSCO

Assembly BUSCO statistics improved after polishing, from 92.0% total complete to 95.9% (Table 2). In the polished, repeat-masked Dovetail assembly, restricting the BUSCO analysis to the largest 10 scaffolds reduced the percentage of duplicated BUSCOs from 7.7% to 3.4% while increasing the percentage of single-copy from 88.2% to 92.6%. Using the Viridiplantae database, the number of total complete BUSCOs is consistent with the result from the assembly of hemp cultivar CBDRx, for which 97% of complete BUSCOs were identified (Grassa *et al.*, 2021). Based on the inclusion of fewer duplicated BUSCO genes in the largest ten scaffolds, we restricted our downstream synteny analyses to the largest ten scaffolds. Figure 1d shows a comparison of the chromosome-level assembly with the PacBio Cascade primary assembly (Padgitt-Cobb *et al.*, 2021), Shinshu Wase assembly (Natsume *et al.*, 2015), and Teamaker assembly (Hill *et al.*, 2017).

**Table 2.** Assembly BUSCO results

| Category | Full assembly or largest 10 scaffolds | Database | % Total complete | % Single-copy complete | % Duplicated complete | % Fragmented | % Missing |
| --- | --- | --- | --- | --- | --- | --- | --- |
| Polished, masked | full | Embryophyta (1614) | 95.9 | 88.2 | 7.7 | 1.3 | 2.8 |
| **Polished, masked** | **10** | **Embryophyta (1614)** | **96.0** | **92.6** | **3.4** | **1.4** | **2.6** |
| Polished, masked | full | Viridiplantae (425) | 97.0 | 90.4 | 6.6 | 0.7 | 2.3 |
| Polished, masked | 10 | Viridiplantae (425) | 97.5 | 94.4 | 3.1 | 0.7 | 1.8 |
| Polished, unmasked | full | Embryophyta (1614) | 94.9 | 87.7 | 7.2 | 1.3 | 3.8 |
| Polished, unmasked | 10 | Embryophyta (1614) | 94.3 | 91.3 | 3.0 | 1.4 | 4.3 |
| Unpolished, unmasked | full | Embryophyta (1614) | 92.0 | 85.5 | 6.5 | 2.3 | 5.7 |

### Genome size and heterozygosity

The haploid size of the genome of *H. lupulus* var. *lupulus* estimated by flow cytometry ranges from 2.57 Gb (Natsume *et al.*, 2015) to 2.989 Gb (Zonneveld *et al.*, 2005) for different cultivars (SI Table S3). Based on a k-mer distribution analysis, we estimated a haploid genome size of 3,058,114,149 bp (3.058 Gb) for Cascade. The genome is ~4.59%-5.47% heterozygous with a read error rate of ~0.48% (SI Table S4). Out of 563,456,691 total DNA short-reads, 561,688,517 DNA short-reads (99.69%) mapped to the Dovetail assembly. The Dovetail assembly is 64.25% composed of repeat sequences, including 62.14% LTRs, 0.19% DNA transposons, 1.76% simple repeats, and 0.03% LINE repeats (Figure 2a).

**Figure 2.**
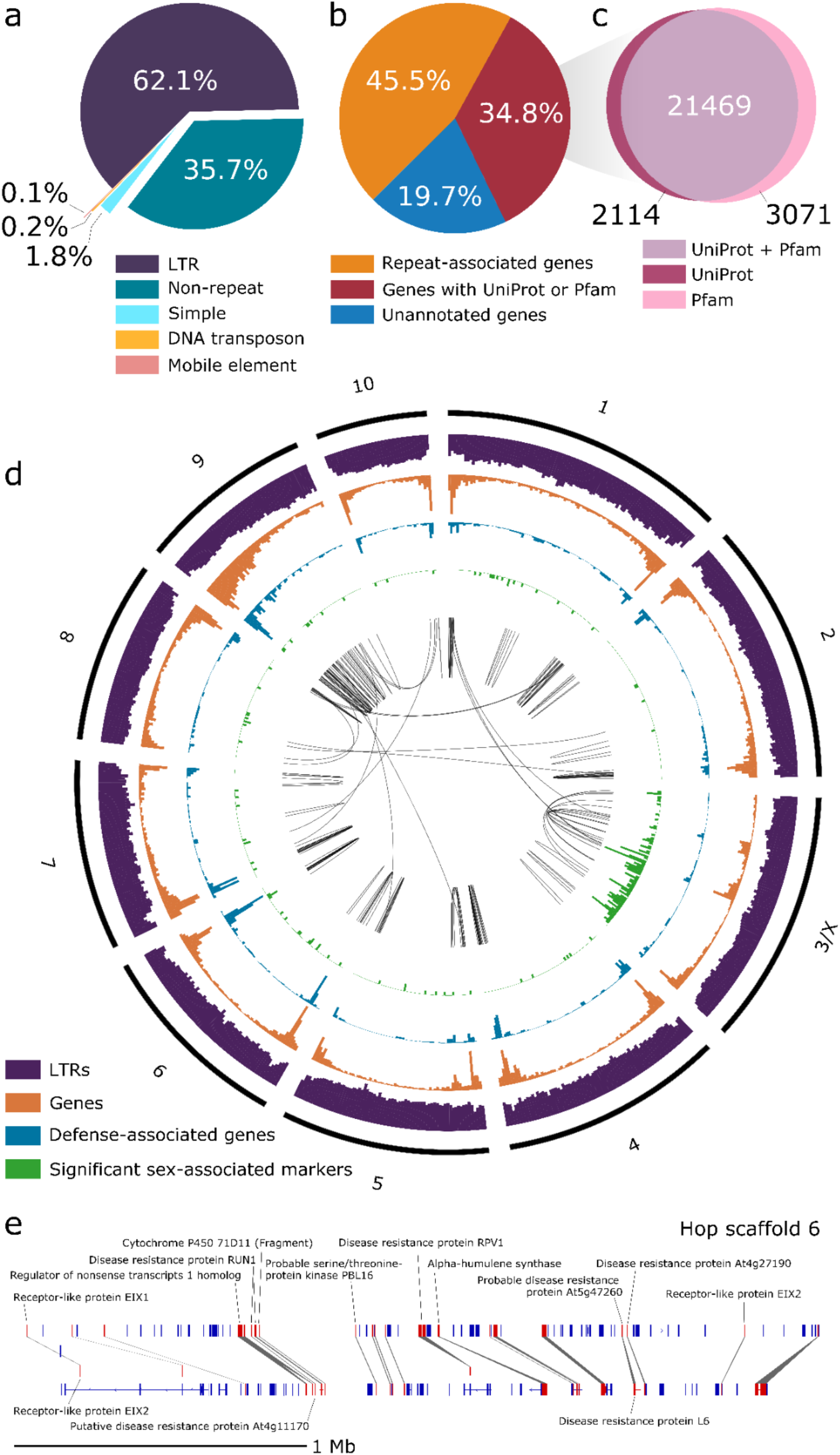
Gene and repeat content in the ten largest assembly scaffolds. **a)** Pie chart showing percentages of different categories of repeat content relative to total repeat content. **b)** Pie chart showing the percentage of genes with similarity to a repeat-associated UniProt protein or Pfam domain, the percentage of genes with similarity to a non-repeat-associated UniProt gene or Pfam domain, and genes lacking similarity to any known UniProt protein or Pfam domain. Venn diagram showing the intersection of genes that have similarity to a UniProt gene and/or a Pfam domain. **d)** Circos plot for the largest ten scaffolds in the Dovetail assembly showing histograms for the density for genes (orange; y-axis range: 4-253), putative defense-response-associated genes (blue; y-axis range: 1-110), long-terminal retrotransposons (purple; y-axis range: 51-5707), and significant sex-associated SNPs (green; y-axis range: 0-12; p-value < 0.05). The center track depicts syntenic blocks within the same scaffold and across different scaffolds. All histograms are split into 5 megabase (Mb) bins, depicting counts per 5 Mb. **e)** Single syntenic block on Dovetail scaffold 6 containing putative disease-response-associated genes and one copy of an alpha-humulene synthase.

### Development of linkage map

The genetic map for mapping population 2017014 resulted in ten linkage groups including a total of 4,090 markers and an overall length of 1,269.5 cM (SI Table S5). Average genetic distance between markers was 0.35 cM with an average of 409 markers per linkage group. Chromosome 7 contained the fewest number of markers (209) while Chromosome 8 had the most (627), reflecting the density of linkage disequilibrium bins formed for each chromosome, as illustrated by average gap size (SI Table S5). SI Figure S5 shows the association between marker positions on the genetic map versus marker location on the physical map for the ten largest scaffolds.

### Gene model content

We estimate 76,595 genes in the hop genome, including 21,698 genes from Transdecoder and 54,897 genes from Maker in the full assembly. For the largest ten scaffolds, this final set includes 34,307 genes from Maker and 20,581 genes from Transdecoder; 94.9% of Transdecoder gene models with expression evidence are on the largest ten scaffolds. Initially, a total set of 71,233 gene models were identified with MAKER (see SI Table S6 for details about Transdecoder gene models). We assessed the completeness of gene models with BUSCO (SI Table S7).

Among the set of 76,595 genes, we identified 34,840 repeat-associated genes and 26,654 genes with similarity to a repeat- and non-repeat-associated gene (Table 3; Figure 2b). We identified 23,583 genes with similarity to a UniProt Embryophyta gene; 20,877 (88.53%) of these genes are present on the ten largest Dovetail scaffolds (Figure 2c). The set of MAKER gene models has fewer gene models with high percent similarity to UniProt genes than Transdecoder (SI Figure S6). The number of genes with GO terms is provided in SI Table S8 (Ashburner *et al.*, 2000). Among the 3,003 genes with putative defense or disease response-associated genes in the full assembly, 2,667 of these genes occur in the largest ten scaffolds.

**Table 3.**
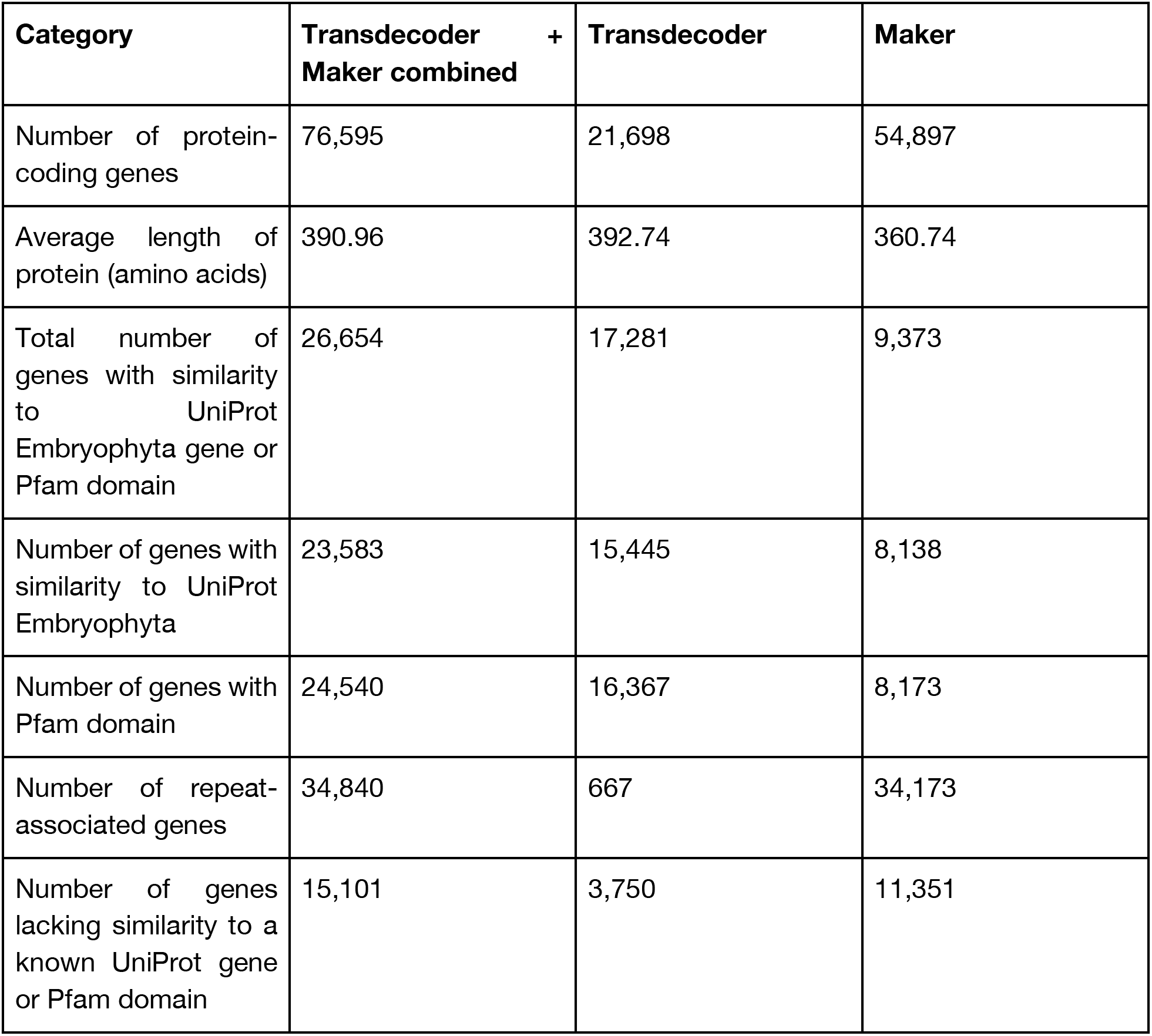
Gene statistics

### Density of genes and long terminal retrotransposons (LTRs)

We visualized the density of genes and LTRs across the ten largest scaffolds in a circos plot (Figure 2d). For most scaffolds, gene density is higher at the ends, which is a pattern observed in other large plant genomes (Dong *et al.*, 2017; Chaw *et al.*, 2019). Defense gene density (blue track) is similar to overall gene density (orange track). The majority of significant sex-associated markers are located on the third largest scaffold (Scaffold_1533; scaffold 3/X), suggesting that scaffold 3 is the X chromosome.

### Analysis of molecular evolution in syntenic gene blocks

Syntenic blocks are depicted in the center track of the circos plot, with most syntenic blocks occurring within the same scaffold (Figure 2d). A depiction of an intra-hop syntenic block on scaffold 6 (Scaffold_172) shows a cluster of disease resistance-associated genes, along with alpha-humulene synthase (Figure 2e). Figure 3a shows syntenic blocks shared between the largest ten scaffolds in the hop and hemp genomes, highlighting extensive sequence similarity shared between hop and hemp. Also highlighted are large regions of genomic sequence that are unique to hop chromosomes, particularly in scaffolds 4, 5, 7, and 10, that potentially correspond to centromeric regions.

**Figure 3.**
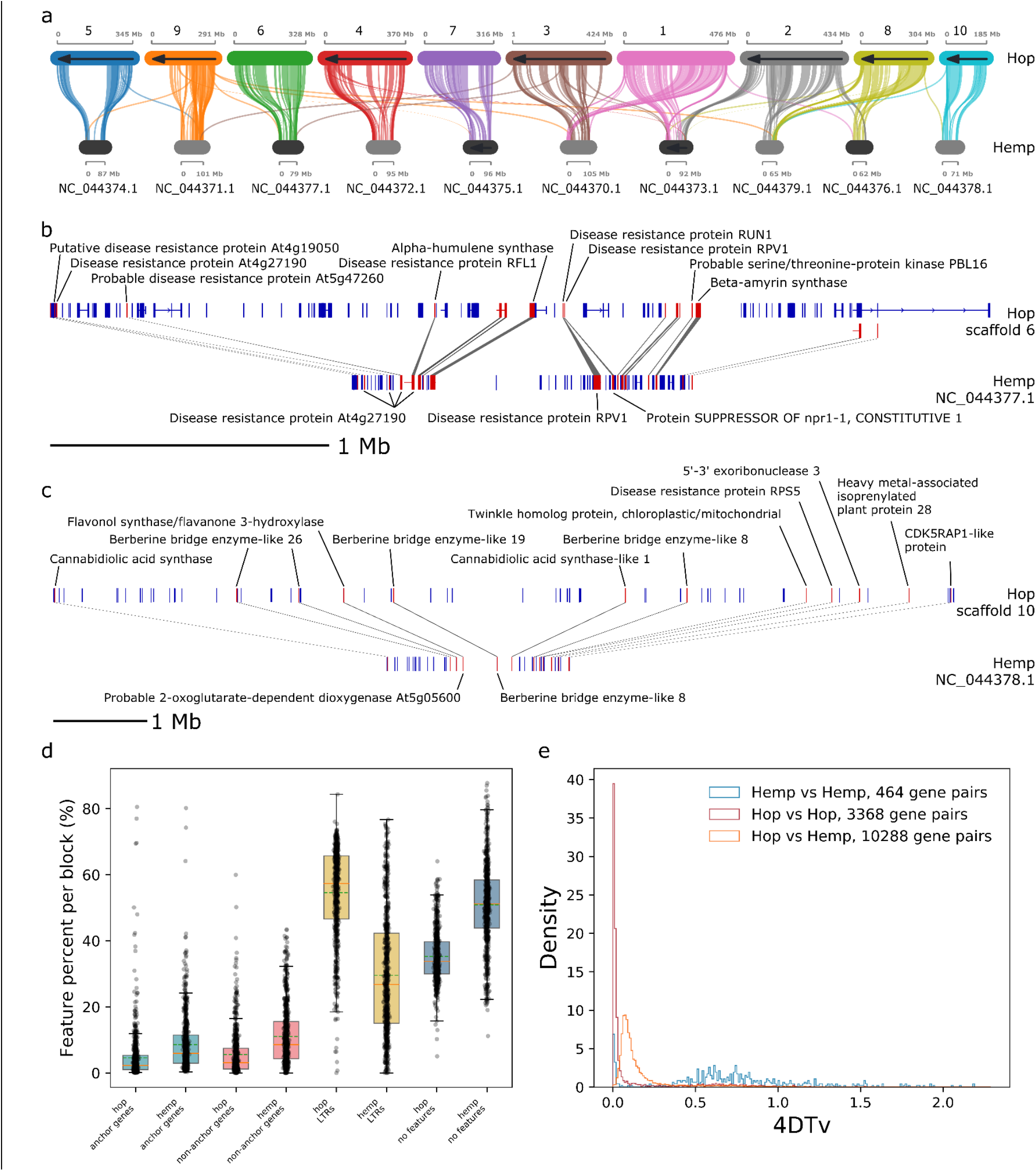
Comparative analysis of hop and hemp syntenic blocks. **a)** Syntenic blocks shared between the largest ten scaffolds in the hop genome assembly and the largest ten scaffolds in the hemp genome assembly. **b)** Single syntenic block between hop scaffold six and hemp scaffold NC_044377.1 containing putative disease-response-associated genes and one copy of an alpha-humulene synthase. **c)** Single syntenic block between hop scaffold ten and hemp scaffold NC_044378.1 containing putative cannabinoid synthesis pathway genes. **d)** Boxplot showing the percentage of syntenic blocks on a genome-wide scale that are composed of anchor genes, non-anchor genes, LTRs per syntenic block, or no annotated feature. **e)** Histogram of 4DTv values from intra-hop, intra-hemp, and inter-hop and hemp.

Figures 3b and 3c show two syntenic blocks shared between hop and hemp. The syntenic block in Figure 3b contains disease response-associated genes and a copy of alpha-humulene synthase. Figure 3c shows genes associated with the cannabinoid synthase pathway, including cannabidiolic acid synthase, cannabidiolic acid synthase-like 1, as well as disease response-associated genes.

On a genome-wide scale, syntenic blocks in hop have expanded LTR content (Figure 3d). The distance between anchor genes in hop is larger than in hemp (SI Figure S7a), and syntenic blocks containing more genes typically have a smaller Ks value, corresponding to more recent large-scale duplication events (SI Figure S7b). We calculated the 4DTv distance (Hellsten *et al.*, 2007) for each collinear anchor gene pair and visualized the distribution to detect large-scale duplication events (Figure 3e) (Vanneste *et al.*, 2015). Among syntenic blocks within hop, and between hop and hemp, enriched GO terms with the greatest statistical significance are associated with energy, metabolism, and development (SI Figures S8 and S9).

### Identification of orthologous genes

A total of 22,739 orthologous gene groups (OGGs) were identified with OrthoFinder (Emms & Kelly, 2019). (SI Figure S10). There are 24,513 hop genes from the ten largest scaffolds in OGGs; 58.3% of OGGs contain hop genes, and 10.5% of hop genes are in species-specific OGGs.

### Estimation of the divergence date using Ks

The optimal number of mixture model components for Ks from hop vs hop anchor genes was three, for hemp vs hemp was two, and for hop vs hemp was three. We interpret the component mean nearest to the modal peak as the primary putative duplication event. For hop vs hop, the component means occur at Ks values of 0.027, 0.251, and 1.616; for hemp vs hemp at Ks values of 0.041 and 1.612; and for hop vs hemp at Ks values of 0.195, 0.329, and 1.605 (Figure 4a).

**Figure 4.**
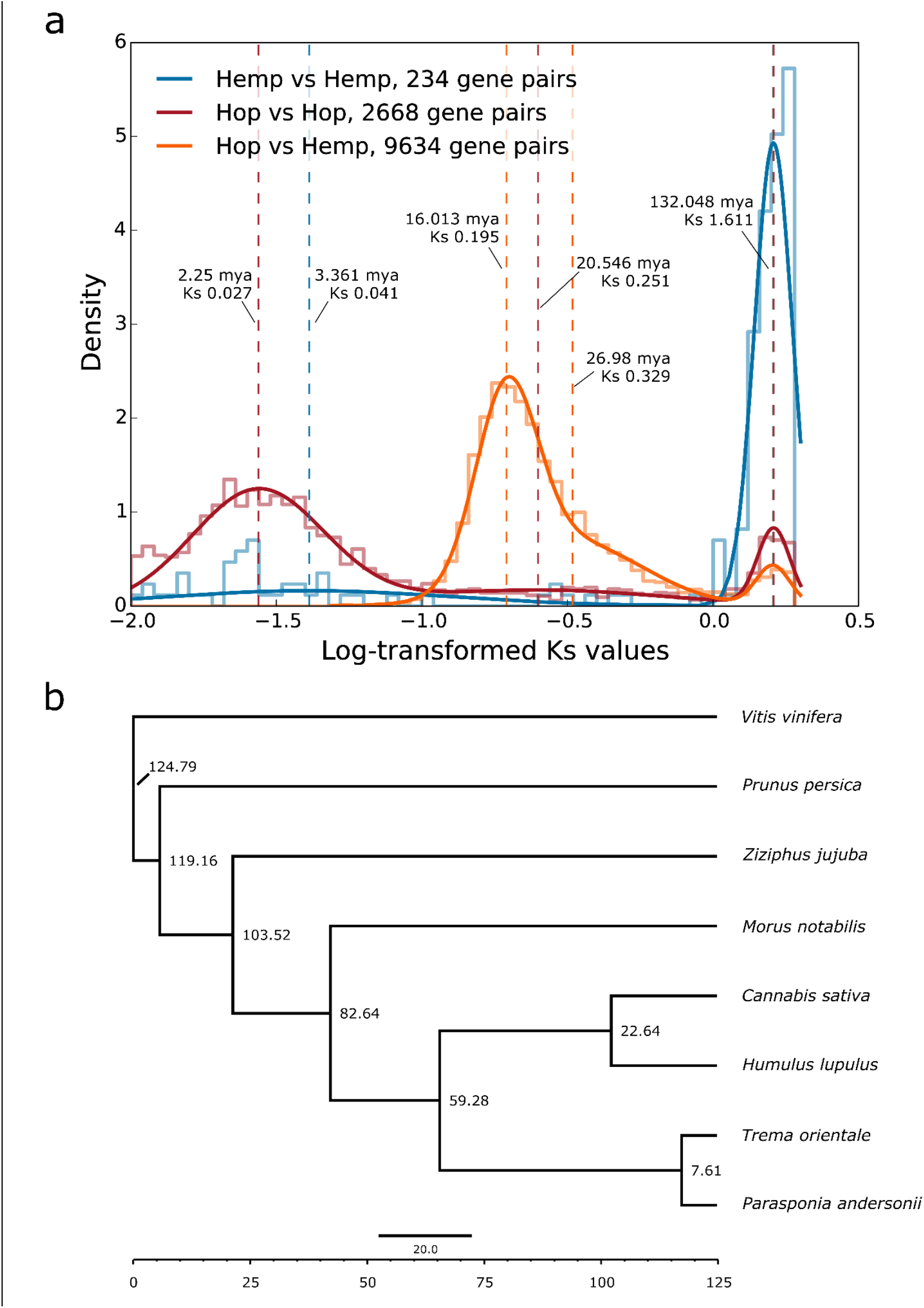
Estimates of divergence time. **a)** Distribution of log-transformed Ks values from collinear gene pairs. The mixture model is superimposed over the histogram, and dashed lines correspond to the location of means identified by the mixture model. Time is included for each dashed line and is calculated using the equation *T* = *Ks*/*2λ* (Koch *et al.*, 2000) and the substitution rate (λ) of 6.1 × 10-9. **b)** Bayesian time tree providing estimated dates of divergence between species. The dates are estimated by MCMCTree, incorporating fossil data and the species tree from OrthoFinder.

Comparing log-Ks and 4DTv distributions, 4DTv shows a strongly-pronounced peak near zero for hop vs hop (Figure 3e), which is less apparent in the Ks distribution because we imposed a minimum Ks threshold of 0.01 (Murakami, 2000). The diffuse peak at 4DTv=0.5-1.0 presumably corresponds to the small, sharp peak in the Ks distribution near Ks=1.611. Overall, both distributions show the same pattern.

For hop vs hemp, the peak occurring at Ks=0.195 putatively marks the primary speciation event. The component mean occurring at Ks=0.329 appears to overlap with the putative primary duplication event at Ks=0.195. We do not necessarily interpret the component mean at Ks=0.329 as a distinct duplication event. The primary putative duplication event shows a positive skew, characteristic of a trend described previously, wherein overfitting in the heavy right tail of the main peak can occur, leading to erroneous detection of duplication events (Zwaenepoel *et al.*, 2019). We calculated a divergence date for hop and hemp of 16.013 mya, based on Ks=0.195, indicating that this λ is concordant with the results of our Bayesian time tree (Figure 4a).

### Fossil-calibrated Time Tree

Based on a larger log-likelihood value, we determined that the independent log-normally distributed relaxed-clock model (clock=2; clock2) (Rannala & Yang, 2007) out-performed the strict molecular clock (clock=1; clock1). The estimated time divergence for *Humulus* and *Cannabis* with the independent log-normally distributed relaxed-clock model is 22.6438 mya (95% highest posterior density [HPD]=15.6728, 28.7994) (Figure 4b).

### Gene family expansion and contraction

We identified contracted and expanded hop gene families (Figures 5a and 5b) and investigated functionally enriched GO terms in expanded and contracted groups (Figures 5c and 5d). There are 571 gene families expanded in both hop and hemp; 3,817 gene families are expanded in hop only and 1,159 gene families are expanded in hemp (Figure 5a). Among contracted gene families, 4,327 gene families are shared between hop and hemp, with 3,296 families specific to hop and 2,118 families specific to hemp (Figure 5b).

**Figure 5.**
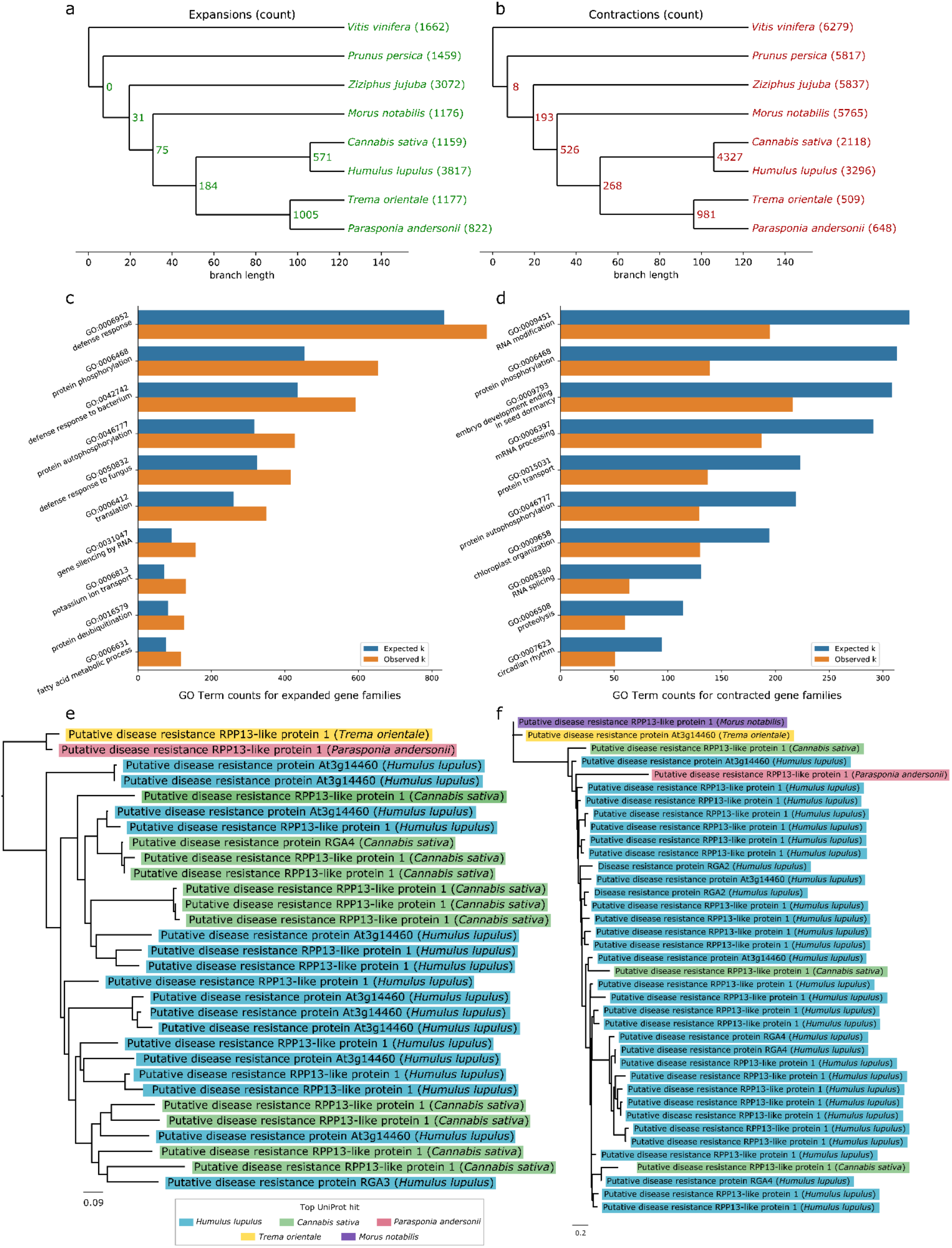
Gene family expansion and contraction. **a)** Tree showing the number of expanded gene families. **b)** Tree showing the number of contracted gene families. **c)** Bar chart showing statistically significant Biological Processes GO terms with the largest number of observed occurrences from expanded families identified with a hypergeometric test. **d)** Bar chart showing statistically significant Biological Processes GO terms with the largest number of observed occurrences from contracted families identified with a hypergeometric test. **e)** Gene family tree containing genes with significant similarity to putative disease resistance genes that are specific to the Cannabaceae family. Each branch is color-coded according to putative functional association. **f)** Gene family tree containing genes with significant similarity to putative disease resistance genes that are specific to the Cannabaceae family, except for one gene that occurs in mulberry (*Morus notabilis*). Each branch is color-coded according to putative functional association.

Among the most significant functionally-enriched Biological Processes GO terms in the expanded gene families are protein phosphorylation (GO:0006468), defense response (GO:0006952, GO:0042742, GO:0050832), and gene silencing by RNA (GO:0031047) (Figure 5c). Among contracted gene families, we find significant functionally enriched Biological Processes GO terms including RNA modification (GO:0009451), protein phosphorylation (GO:0006468), embryo development ending in seed dormancy (GO:0009793), and circadian rhythm (GO:0007623) (Figure 5d). We further highlight two gene families associated with Biological Processes GO term “defense response” containing putative disease resistance genes that are expanded in hop (Figures 5e and 5f), and are also mostly restricted to the Cannabaceae.

### Orthologous gene groups containing genes associated with terpene and cannabinoid biosynthesis

We highlight two phylogenetic trees in Figure 6 featuring secondary metabolic pathways of interest. Figure 6a depicts a tree containing genes with similarity to terpene synthases. Further experimental validation is needed to confirm their functional activity, as sequence similarity is not sufficient to assign function. Figure 6b shows the orthologous gene group containing cannabidiolic acid synthase, cannabidiolic acid synthase-like 1 and 2, tetrahydrocannabinolic acid synthase, inactive tetrahydrocannabinolic acid synthase, and Berberine bridge enzyme-like genes. The full-length CDS in hop (HUMLU_CAS0069948.t1.p1) with similarity to cannabidiolic acid synthase is expressed, and is most similar to a gene in *Cannabis* (XP_030502671.1) that is annotated as cannabidiolic acid synthase in our annotation, and a cannabidiolic acid synthase-like gene in the annotation of the hemp CBDRx assembly (Grassa *et al.*, 2021). The only expressed copy of cannabidiolic acid synthase in hemp (XP_030480746.1) is present in the gene tree in Figure 6b, clustering near copies of cannabidiolic acid synthase-like 1, Inactive tetrahydrocannabinolic acid synthase, and a copy of cannabidiolic acid synthase-like 1 in hop (HUMLU_CAS0071292.t1.p1).

**Figure 6.**
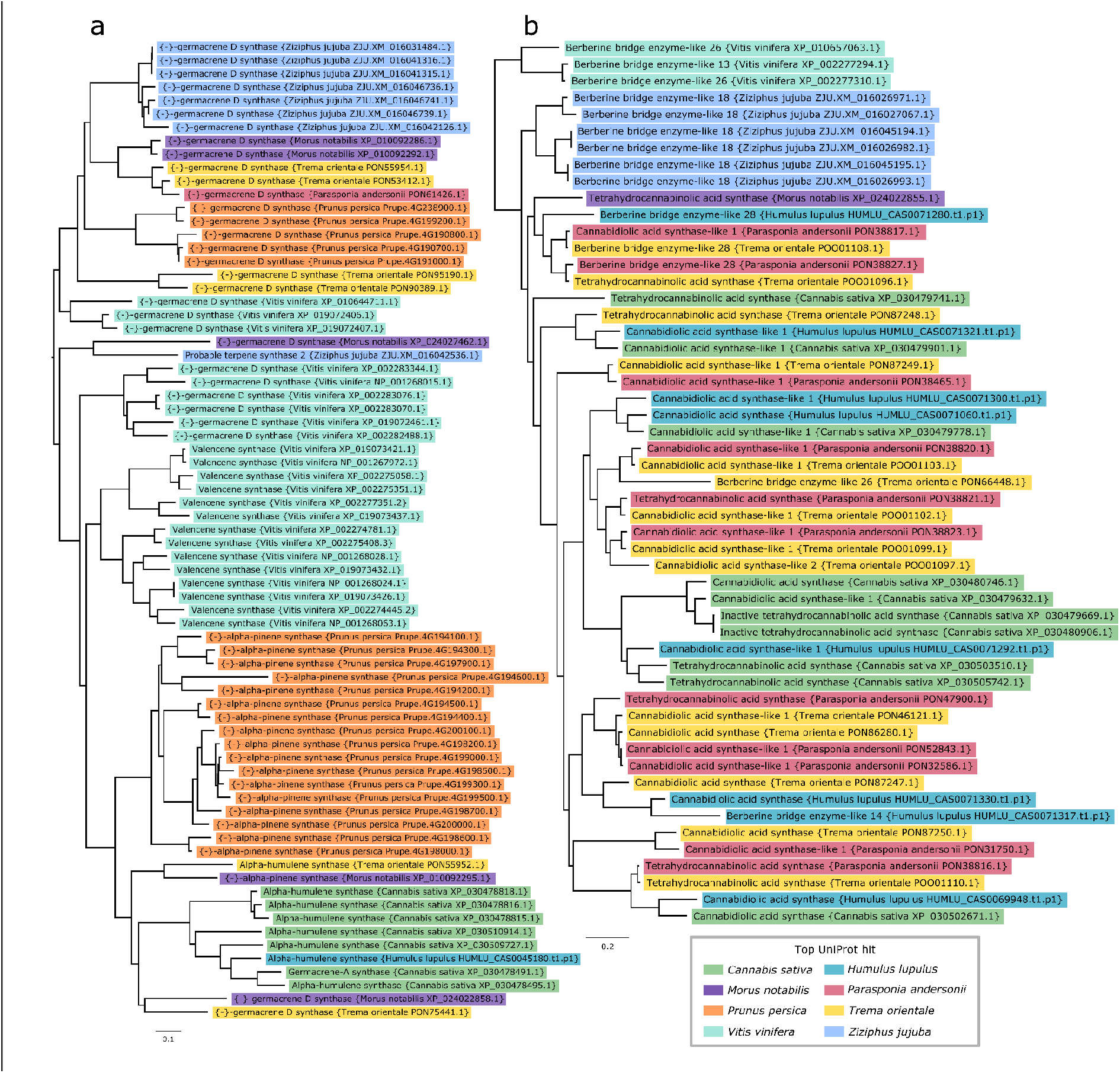
Gene family trees associated with terpene and cannabinoid biosynthesis. **a)** Gene family tree containing terpene synthase genes that is color-coded according to putative functional association. **b)** Gene family tree containing genes putatively associated with cannabinoid synthase genes or Berberine-bridge enzyme-like genes. Each branch is color-coded according to putative functional association.

## Discussion

### Genome assembly analysis

We have presented a chromosome-level assembly of Cascade that contains 3.47 Gb of genomic sequence in the largest ten scaffolds, which we expect corresponds to the ten chromosomes. The haploid size of the hop genome was previously estimated to be between 2.57 Gb (Natsume *et al.*, 2015) and 2.989 Gb (Zonneveld *et al.*, 2005) (SI Table S3), and our k-mer-based estimate of the size of the Cascade hop genome is closer to the size of the assembly at 3.058 Gb. The size of the Dovetail assembly is larger than the estimated genome size by flow cytometry; however, the reported genome sizes in the literature demonstrate variation in genome size across cultivars. Variation in genome size is known to occur within plants of the same species (Ohri, 1998) as a result of large structural variants (Saxena *et al.*, 2014).

Cascade has a highly heterozygous genome, which can be desirable in the cultivation of new varieties (Henning *et al.*, 2004). Using short-read DNA sequencing, we estimated the heterozygosity of the hop genome to be approximately 5%, which is similar to the range of other heterozygous genomes, including potato (4.8%) (Leisner *et al.*, 2018), *Vitis vinifera* cultivar Börner (3.1%) (Holtgräwe *et al.*, 2020), *Vitis vinifera* cultivar PN40024 (7%) (Jaillon *et al.*, 2007), and sunflower (10%) (Hübner *et al.*, 2019). Heterozygosity does present challenges in assembly and annotation related to distinguishing between haplotype and paralogous sequences, especially in the case of recent duplications. Although we performed phasing and further efforts to detect haplotype contigs in the PacBio long-read assembly that provided the basis for the Dovetail assembly, heterozygosity continues to be a challenge for future work to overcome (Padgitt-Cobb *et al.*, 2021). Hi-C construction of scaffolds does appear to reduce the inclusion of homologous primary contigs representing the corresponding haplotype. Further restricting BUSCO analysis to the largest ten scaffolds reduces the number of duplicated BUSCOs (7.7% to 3.4%) while increasing the number of single-copy BUSCOs (88.2% to 92.6%), supporting the inclusion of the ten scaffolds for downstream analyses that are sensitive to duplication. Polishing of the assembly using short-read Illumina sequencing further improved the quality of the scaffold sequences, based on the recovery of BUSCO genes.

Zhang *et al.* showed obvious segregation distortions in genetic mapping of hop, presumably caused by multivalent formation during meiosis (Zhang *et al.*, 2017). These segregation distortions could ultimately influence marker positioning on linkage groups with the consequence of large-scale marker misplacement on genetic maps. The genetic map developed for our mapping population, USDA 2017014, was based upon SNP markers identified by genotyping-by-sequencing (GBS) of 281 offspring. Comparisons between the physical position of SNPs and the genetic map (SI Figure S5) show marked divergences between the physical and genetic positions. Our results demonstrate the potential problems posed by using genetic maps to assemble contigs into chromosomal-scale scaffolds. Long-range interaction-based methods such as Dovetail Hi-C, coupled with PacBio long-read sequencing, allow for the assembly of contigs into chromosomal-scale scaffolds independently of genetic maps. Based upon our results it is recommended that future hop genome assemblies avoid use of genetic maps for scaffolding of large contigs and instead use methods such as long-range scaffolding with Hi-C or more traditional large insert bacterial artificial chromosome (BAC) libraries that overlap genomic regions.

Our new estimate of repeat content of the Dovetail assembly for Cascade, at 64.25%, is similar to previous estimates for repeat content, with repeat content for *H. lupulus* Japanese wild hops at 60.1%, var. *lupulus* at 61.3%, and var. *cordifolius* at 59.2% (Pisupati *et al.*, 2018). Previously, in the PacBio long-read assembly, we estimated the total repeat content for the assembly at 71.46% (Padgitt-Cobb *et al.*, 2021). Assembly with Hi-Rise did result in 1,027 breakage points to the PacBio assembly, as well as 8,131 joins, and it is possible that these breakages resulted in a shuffling of genome content that changed the resulting total repeat percentage. An earlier estimate of repeat content for *H. lupulus* cultivar Shinshu Wase was 34.7% (Natsume *et al.*, 2015). However, assemblies generated with short-read sequencing likely underestimate repeat content by not comprehensively capturing intergenic and repeat regions. The repeat content of the closely-related *C. sativa* genome is 64% (Pisupati *et al.*, 2018). Although we observe syntenic blocks expanded in hop due to LTR insertion, on a genome-wide scale, the similarity in repeat content between hop and hemp suggests that LTRs do not contribute to larger genome size in hop.

### Genomic content of syntenic blocks

The syntenic blocks in Figures 3b and 3c show an expansion of LTR sequences in hop, and we observe on a genome-wide scale that syntenic blocks in hop have expanded LTR content (Figure 3d). LTRs impact gene expression by altering the spacing and organization of accessible chromatin regions (ACRs), which can be involved in regulation of gene expression by harboring accessible *cis*-regulatory elements (CREs), including transcription factor binding sites (Zhao *et al.*, 2018). In maize, single-nucleotide polymorphisms in ACRs are responsible for ~40% of heritable variance in quantitative traits, highlighting the importance of identifying ACRs containing regulatory DNA (Rodgers-Melnick *et al.*, 2016). For hop, an open question is whether the apparent expansion of LTR content in syntenic blocks has influenced the evolution and regulation of genes involved in traits of interest. Future work will be necessary to identify CREs, ACRs, and assess their role in controlling gene expression (Jensen *et al.*, 2021).

### Gene family expansion and contraction

Among the most enriched GO terms in both expanded and contracted gene families, the GO term “protein phosphorylation” (GO:0006468) occurs as a significant Biological Processes GO term for both contracted and expanded gene families. Hop genes with similarity to UniProt genes associated with protein phosphorylation include Disease resistance protein RPP8 (Q8W4J9), chloroplastic Geranylgeranyl diphosphate reductase (Q9ZS34), Protein PHYTOCHROME-DEPENDENT LATE-FLOWERING (F4IDB2), Ultraviolet-B receptor UVR8 (Q9FN03), LEAF RUST 10 DISEASE-RESISTANCE LOCUS RECEPTOR-LIKE PROTEIN KINASE-like 1.2 (P0C5E2), and Two-component response regulator-like APRR1 (Q9LKL2), which is involved in light-mediated flowering response (Matsushika *et al.*, 2000). Although we observe genes with the same putative function occurring in both expanded and contracted gene families, in our analysis these genes cluster into different orthologous groups, suggesting genes with these putative functions have undergone duplication and sub-functionalization (Panchy *et al.*, 2016).

The trees in Figures 5e and 5f contain disease response-associated genes present in the expanded gene families. Both gene trees contain multiple copies of RPP13-like protein 1, which is associated with resistance to downy mildew in *Arabidopsis*, and is characterized by high amino acid divergence within the functional domain (Bittner-Eddy *et al.*, 2000). Resistance gene analogs (RGA2, RGA3, RGA4) are also present in the two trees (Sekhwal *et al.*, 2015).

### Date of species divergence

We estimate the divergence date for hop and hemp to be approximately 16.013 mya using Ks (Figure 4a) and approximately 22.64 mya with the Bayesian inference-based time tree. Our new estimates of time divergence for hop and hemp approximately agree with previous estimates between 18.23 and 27.8 mya (Zerega *et al.*, 2005; Divashuk *et al.*, 2014; McPartland, 2018; Zhang *et al.*, 2018a; McPartland *et al.*, 2019; Kovalchuk *et al.*, 2020; Jin *et al.*, 2020). Each of these estimates is accompanied by its own uncertainty interval, increasing the overall range of divergence time. Our specific point estimates for the divergence time fall within a narrower and slightly more-recent window than previous estimates; however, if we take into account the HPD interval of 15.6728 to 28.7994 from the Bayesian time tree, our results are consistent with previous estimates.

The time equation, *T* = *Ks*/*2λ*, is dependent on the value of λ, and there are different estimates of λ among plants, including 6.1 × 10^−9^ (Lynch & Conery, 2000) and 1.5 × 10^−8^ (Koch *et al.*, 2000) for *Arabidopsis*, and a range of 2.1 × 10^−9^ to 2.9 × 10^−9^ for the monocot-dicot divergence (Wolfe *et al.*, 1987). If we apply other values of λ, including λ=2.1 × 10^−9^ (Wolfe *et al.*, 1987; Murakami, 2000), we calculate a divergence date of 46.43 mya. With λ=1.23 × 10^−9)^ (Xiang *et al.*, 2000), we calculate a divergence date of 79.26 mya, which is closer to the crown age of Cannabaceae, estimated to be 87.4 mya (Jin *et al.*, 2020). Given that our estimated divergence date using λ=6.1 × 10^−9^ agrees with our date estimated by Bayesian inference, along with its application to closely-related species hemp and mulberry (He *et al.*, 2013), suggests that it is reflective of the rate of evolution in hop.

The overall sparsity of the fossil record, as well as changes to the assignment of fossils, speaks to the uncertainty of the results. Multiple fossils from extinct *Humulus* species are dated to the Oligocene (23.03-33.9 mya) (McPartland, 2018). Older, more debatable fossil evidence for *Humulus* includes a leaf fossil, with a date of 34.07 mya based on ^40^Ar/^39^Ar radiometric dating (Meyer, 2003; McPartland, 2018), from Florissant, Colorado, USA (MacGinitie, 1953). However, this older fossil remains questionable because it was originally identified as *Vitis* (MacGinitie, 1953), and subsequently re-assigned to *Humulus* (MacGinitie, 1969). Collinson estimated the time of origin for *Humulus* and the extinct genus *Humularia* at the boundary of the Eocene and Oligocene epochs, corresponding to 33.9 mya (Collinson, 1989); however, this date hinges on the reliability of the Florissant leaf fossil, which McPartland notes was insufficient evidence to warrant conclusive assignment to either family based on the lack of diagnostic fruit (Boutain, 2014; McPartland, 2018). Based on the uncertainty related to placing this fossil, we opted not to incorporate it into our fossil calibration.

### Conclusion and future work

Our chromosome-level genome assembly of Cascade lays the foundation for further evolutionary and biodiversity studies focusing on *Humulus* and the Cannabaceae. This genomic resource will provide a better understanding of content and organization of genes involved in flowering time (Salvi *et al.*, 2007), growth and development, defense response, and metabolism. Future work is needed to identify and map centromeres and telomeres of the hop genome, and to resolve biodiversity and structural variation across *Humulus lupulus* cultivars.

## Acknowledgements

We thank Dovetail Genomics for Hi-C sequencing and assembly. We thank Chris Sullivan and Kenneth Lett at the Center for Quantitative Life Sciences (CQLS) at Oregon State University for assistance with the computing infrastructure. This work was supported by USDA-ARS CRIS Project 5358-21000-040-00D and Hopsteiner. LPC is supported by an AFRI Predoctoral Fellowship (grant no. 2020-67034-31722) from the USDA National Institute of Food and Agriculture.

## Author Contribution

LPC, PM, JH, and DH designed the study. LPC polished and annotated the genome, analyzed and interpreted data, created figures, and wrote the manuscript. NP prepared RNA-seq samples and contributed text about RNA-seq preparation. DH analyzed and interpreted data, and provided guidance about figure design and significant edits to the manuscript. JH developed the mapping population and genetic map, and developed the associated figures and manuscript sections for the mapping population and genetic map. LPC, PM, JH, and DH reviewed and edited the manuscript. JH and PM acquired funds to perform the study.

## Data Availability Statement

The data that support the findings of this study are openly available on the Downloads page of http://hopbase.cqls.oregonstate.edu/ and under NCBI BioProject ID PRJNA562558. Analysis pipelines, scripts, and specific commands are included on the GitHub project page, at https://github.com/padgittl/CascadeDovetail.

