## Supplementary Information for "A chromosome-level assembly of the “Cascade” hop (Humulus lupulus) genome uncovers signatures of molecular evolution and improves time of divergence estimates for the Cannabaceae family"

The following Supporting Information is available for this article:

**Fig. S1.** Distribution of Dovetail Hi-C read insert sizes

**Fig. S2.** Dovetail Hi-C link density histogram

**Fig. S3.** Plot showing contiguity of the input assembly and the final HiRise scaffolds

**Fig. S4.** Nucleotide, dinucleotide, and trinucleotide content of the assembly

**Fig. S5.** Genetic map vs physical map positions

**Fig. S6.** Percent identity to UniProt genes among Transdecoder and MAKER gene models

**Fig. S7.** Comparison of inter-anchor distances and average Ks values in hop and hemp syntenic blocks

**Fig. S8.** Functional enrichment of GO terms in hop vs hop syntenic blocks

**Fig. S9.** Functional enrichment of GO terms in hop vs hemp syntenic blocks

**Fig. S10.** OrthoFinder results

**Fig. S11.** MCMCTree convergence

**Table S1.** Statistics about Hi-C libraries and HiRise assembly

**Table S2.** Analysis of polished assembly quality

**Table S3.** Estimated genome sizes of *Humulus* and closely-related species

**Table S4.** Hop genome heterozygosity and repeat content based on short-read DNA sequencing

**Table S5.** Linkage group statistics for the mapping population USDA 2017014

**Table S6.** Transdecoder gene model results.

**Table S7.** Gene model BUSCO results

**Table S8.** Number of genes with GO terms

**Table S9.** Pfam repeat-associated domains (this table is provided as a separate Excel file)

**Table S10.** Conditional repeat Pfam domains

**Methods S1** Detailed descriptions of SNP identification and filtering; genome size and heterozygosity; gene model development, quality assessment, and assignment of putative function; and molecular evolutionary analyses.

**Supporting References**

**Fig. S1. Distribution of Dovetail Hi-C read insert sizes.** Scatter plot showing the distribution of insert sizes in the Dovetail library. The distance between the forward and reverse reads is given on the x-axis in base pairs, and the probability of observing a read pair with a given insert size is shown on the y-axis.

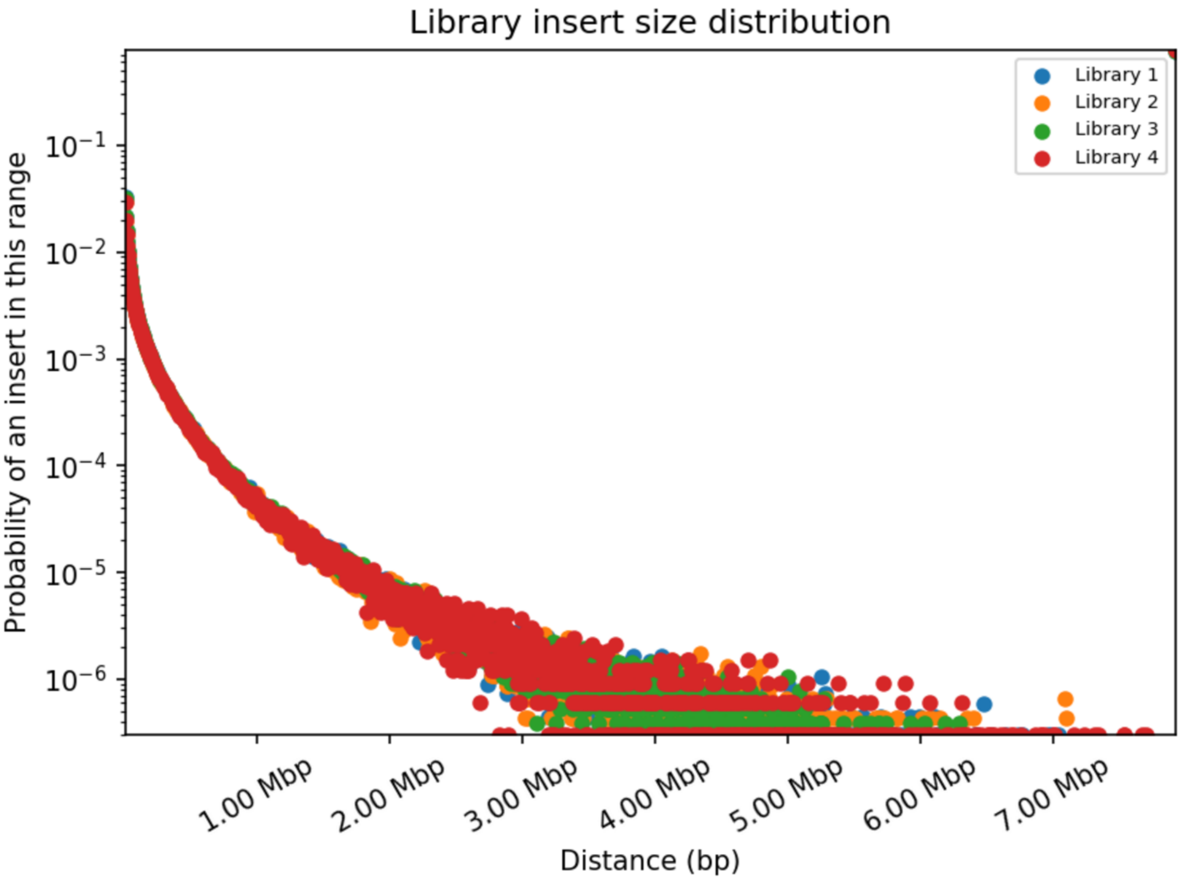

**Fig. S2. Dovetail Hi-C link density histogram.** This link density histogram shows the mapping positions of the first and second read in the read pair respectively, grouped into bins on the x- and y-axes. The color of each square gives the number of read pairs within that bin. White vertical and black horizontal lines have been added to show the borders between scaffolds. Scaffolds less than 1 Mb are excluded.

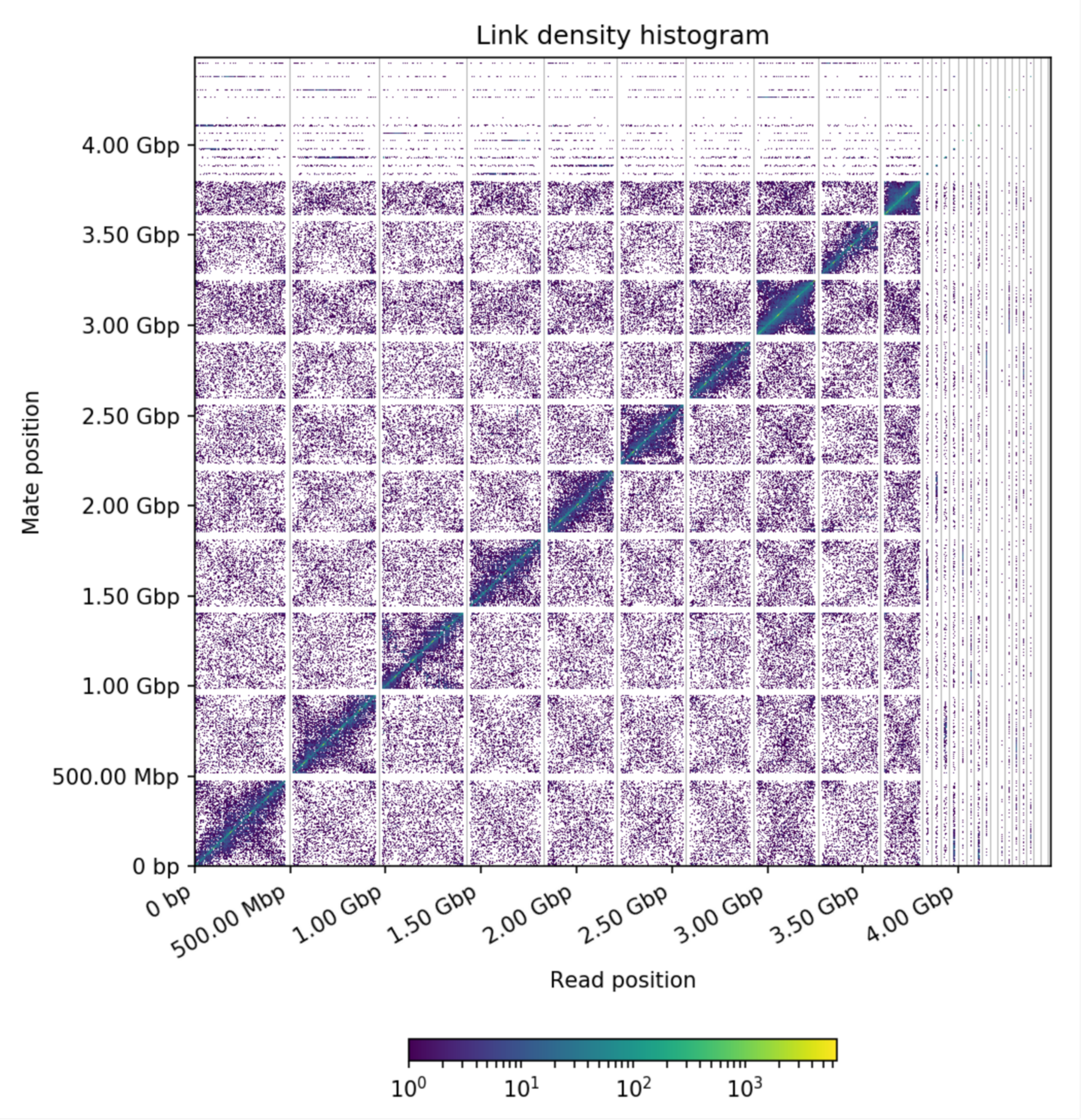

**Fig. S3. Plot showing contiguity of the input assembly and the final HiRise scaffolds.** This plot shows a comparison of the contiguity of the input assembly and the final HiRise scaffolds. Each curve shows the fraction of the total length of the assembly present in scaffolds of a given length or smaller. The fraction of the assembly is indicated on the y-axis and the scaffold length in base pairs is given on the x-axis. The two dashed lines mark the N50 and N90 lengths of each assembly.

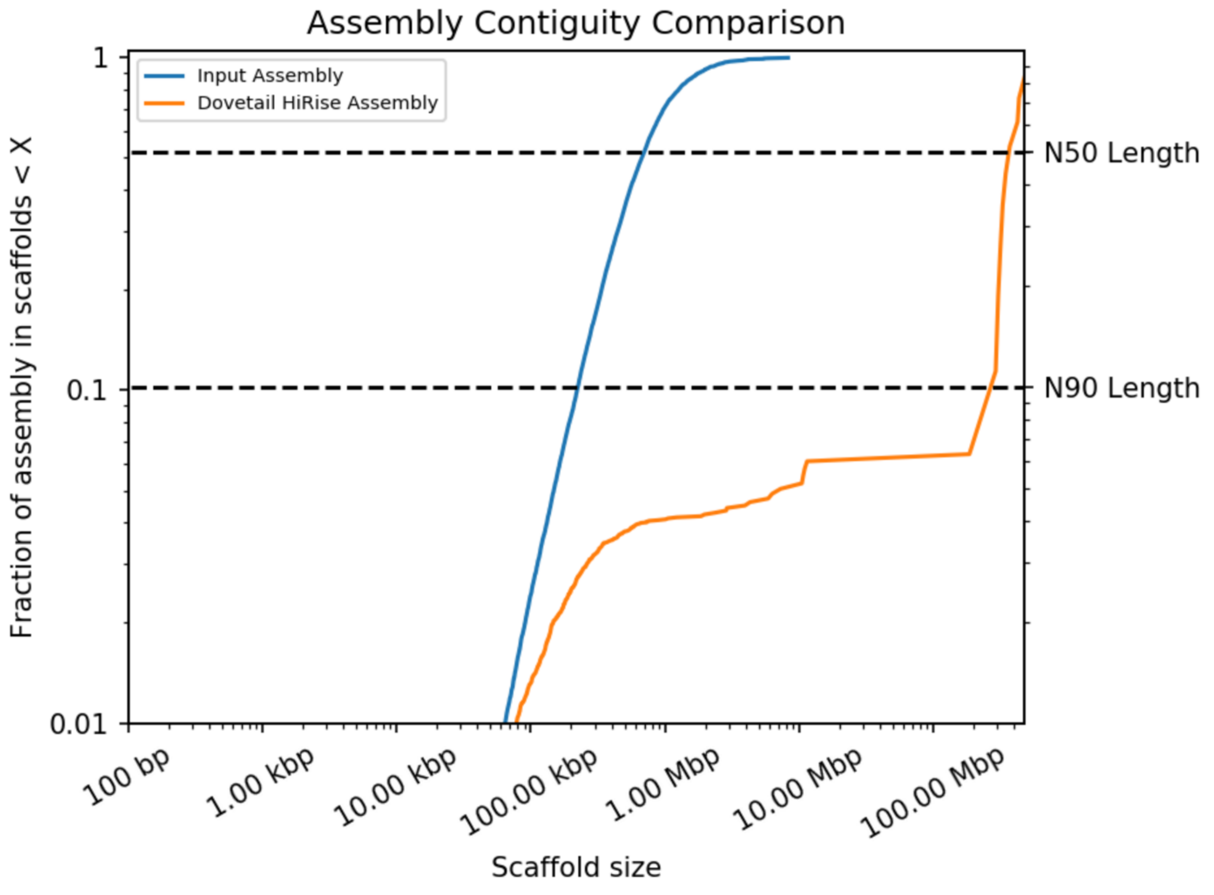

**Fig. S4. Nucleotide, dinucleotide, and trinucleotide content of the assembly. a)** A pie chart shows the percentages for each nucleotide in the largest ten scaffolds. **b)** Heatmap for the dinucleotide composition showing the enrichment of a given dinucleotide. The enrichment score is calculated by dividing the observed frequency of a dinucleotide by the expected frequency. The dinucleotide CG shows depletion, with an enrichment score of 0.66. **c)** Scatter plot showing the expected frequency of a trinucleotide on the x-axis along with the observed frequency on the y-axis. The trinucleotide CHH in particular occurs more frequently than expected; CHH is associated with DNA methylation.

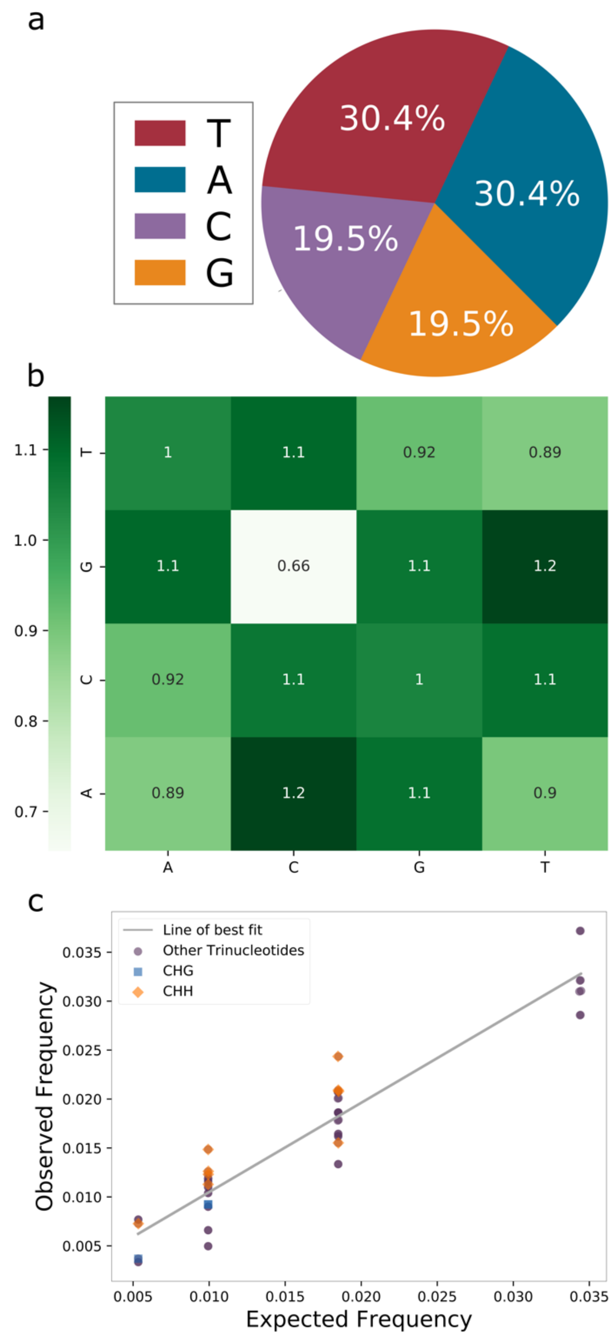

**Fig. S5. Genetic map vs physical map positions.** Comparison between marker positions on the genetic map versus actual marker location on the physical map for the 10 largest scaffolds. Genetic map units are based upon centimorgans (cM) while the physical map simply shows relative position along the scaffold. Positions on the physical map are normalized for comparison to the genetic map.

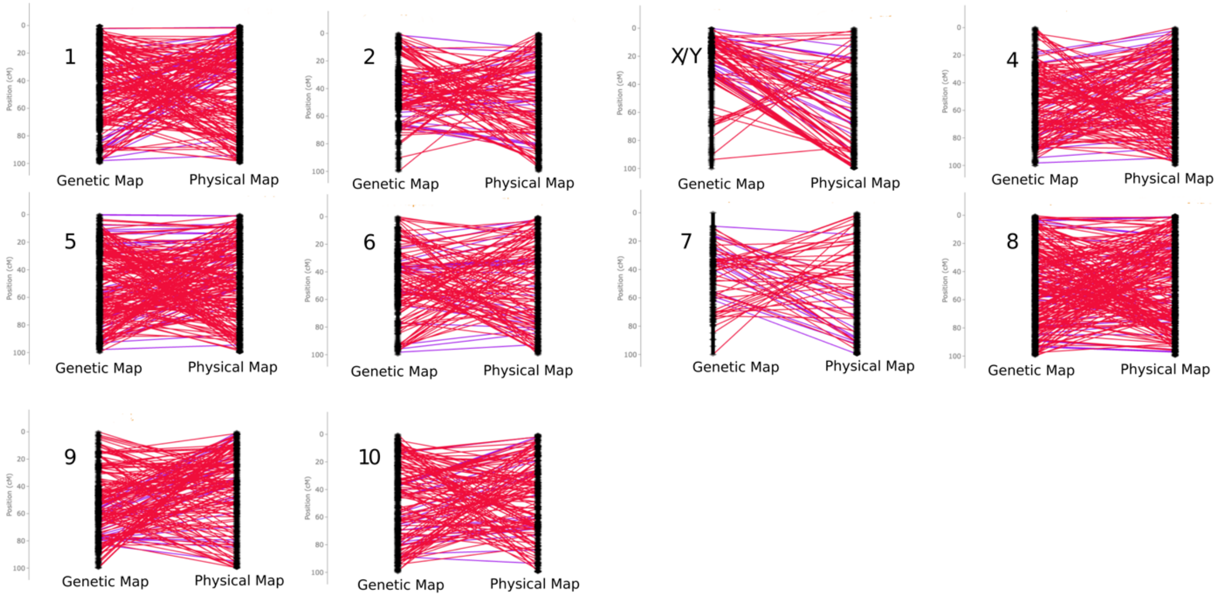

**Fig. S6. Percent identity to UniProt genes among Transdecoder and MAKER gene models.** Histogram of the percent identity to UniProt genes among the set of MAKER gene models and the set of Transdecoder gene models. Transdecoder gene models overall share greater similarity with known genes than Maker gene models.

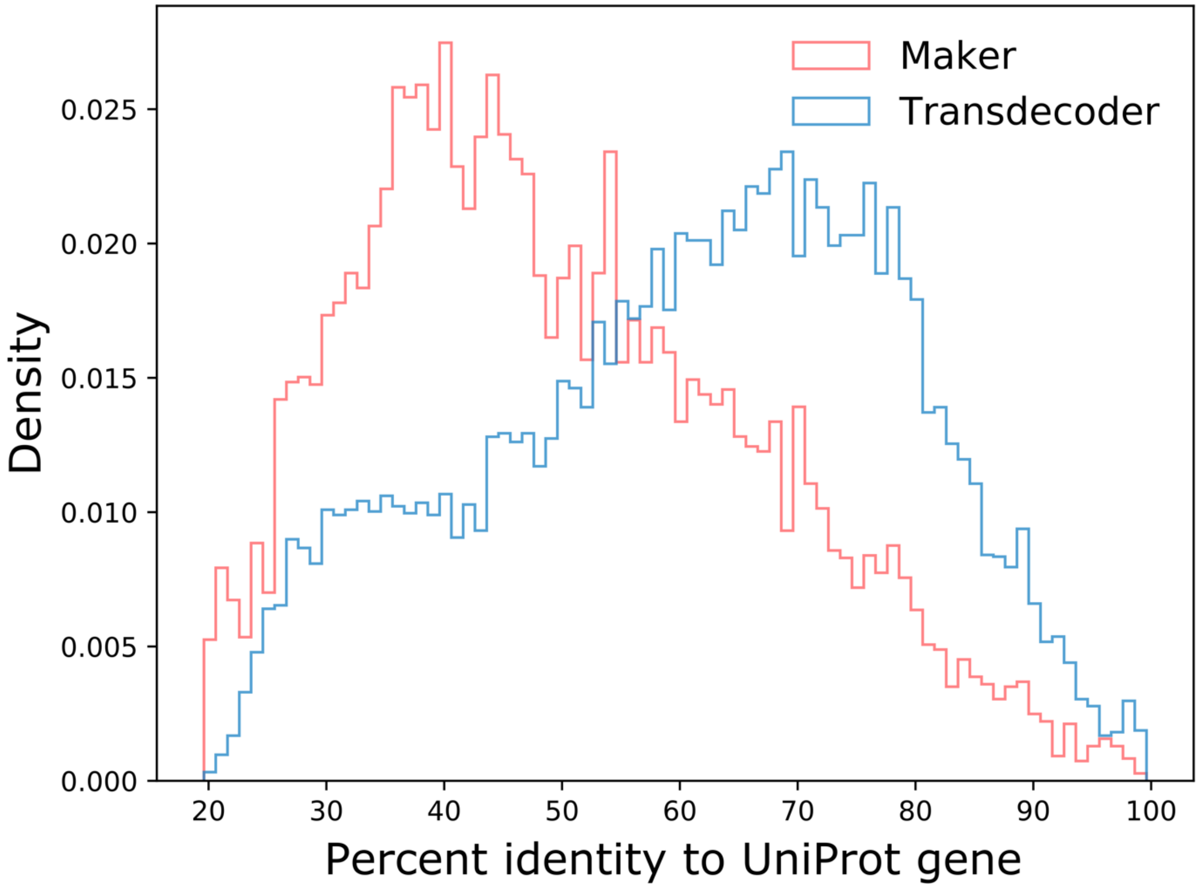

**Fig. S7. Comparison of inter-anchor distances and average Ks values in hop and hemp syntenic blocks. a)** This scatter plot shows the total inter-anchor distance for each syntenic block in hop syntenic blocks and hemp syntenic blocks, corresponding to the distance between anchor genes. **b)** This scatter plot shows the average Ks value per syntenic block on the x-axis and the number of genes in the corresponding syntenic block on the y-axis.

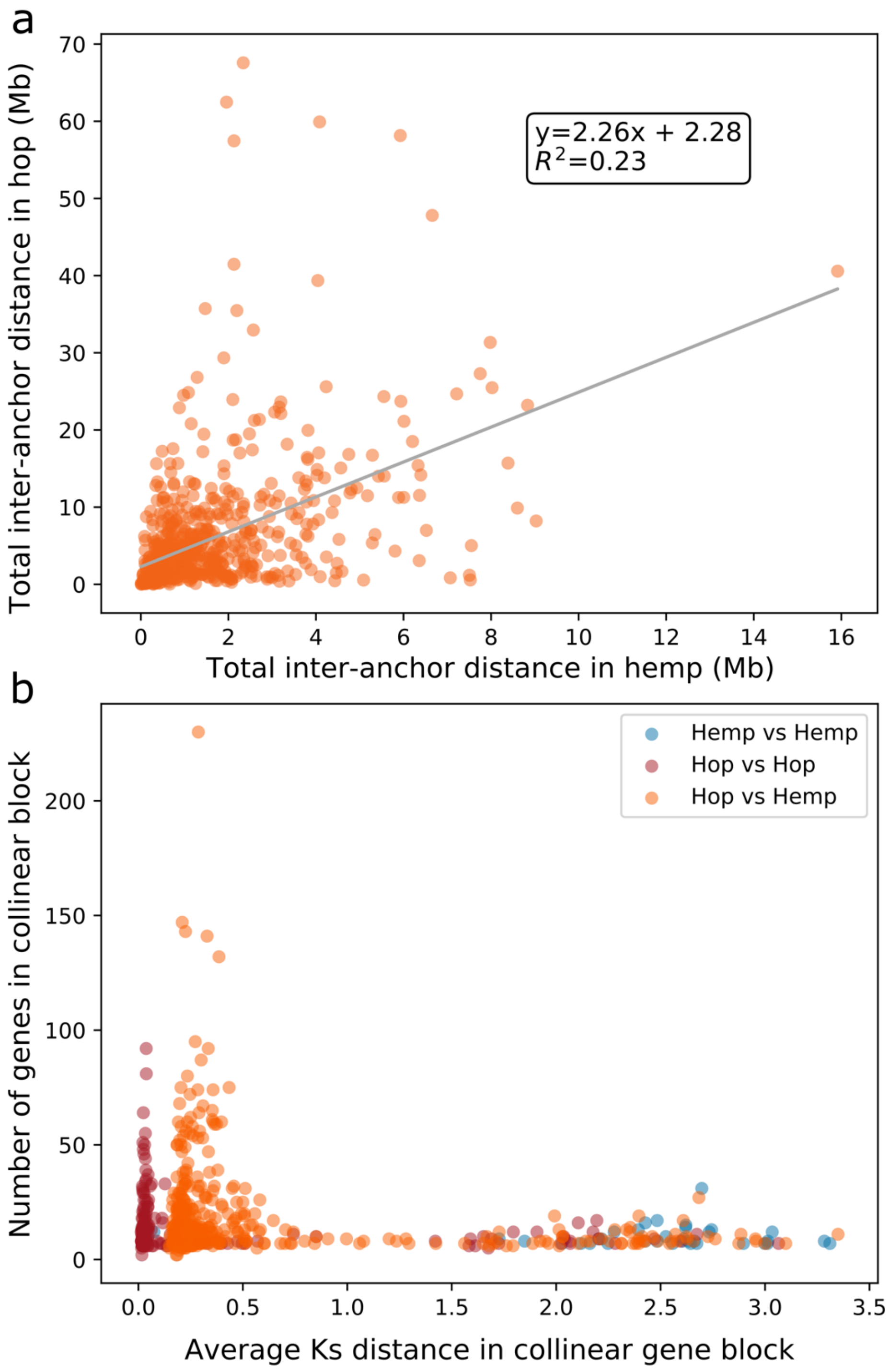

**Fig. S8. Functional enrichment of GO terms in hop vs hop syntenic blocks.** This scatter plot shows functional enrichment and depletion for hop genes in syntenic gene blocks present within the largest ten scaffolds in hop. The labeled GO terms are among the most statistically significant GO terms that have an observed count of at least six. The gradient for the color bar is shaded according to Q-value. Functional enrichment for **a)** biological processes; **b)** cellular components; and **c)** molecular function.

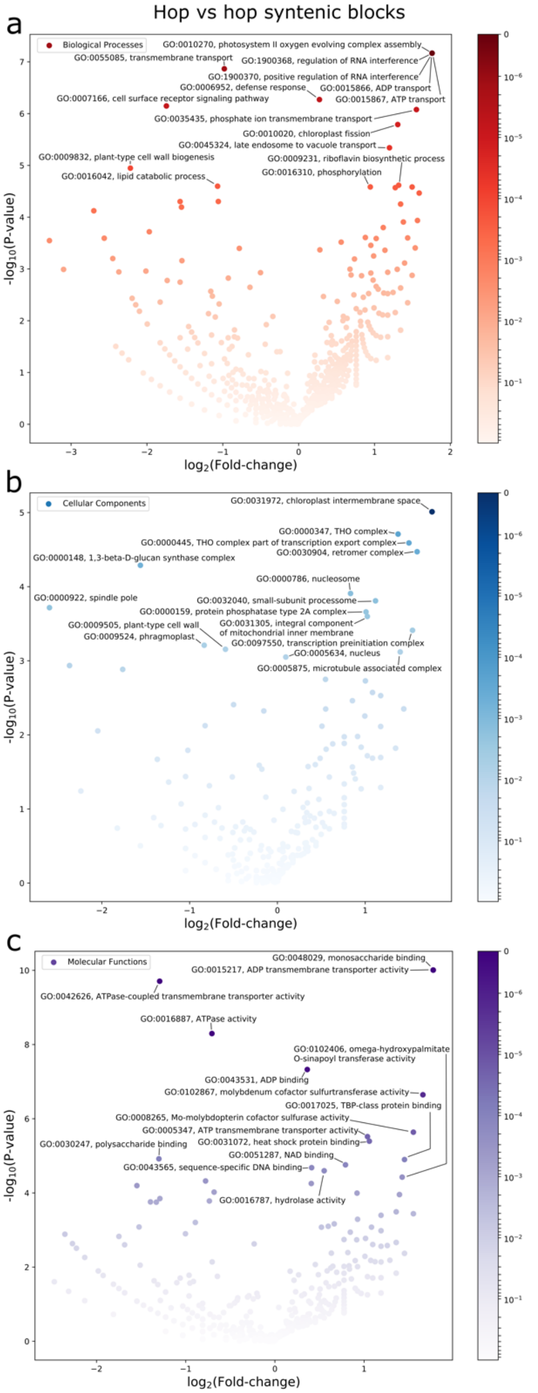

**Fig. S9. Functional enrichment of GO terms in hop vs hemp syntenic blocks.** This scatter plot shows functional enrichment and depletion for hop genes in syntenic gene blocks shared between the largest ten scaffolds in hop and largest ten scaffolds in hemp. The labeled GO terms are among the most statistically significant GO terms that have an observed count of at least six. The gradient for the color bar is shaded according to Q-value. Functional enrichment for **a)** biological processes; **b)** cellular components; and **c)** molecular function.

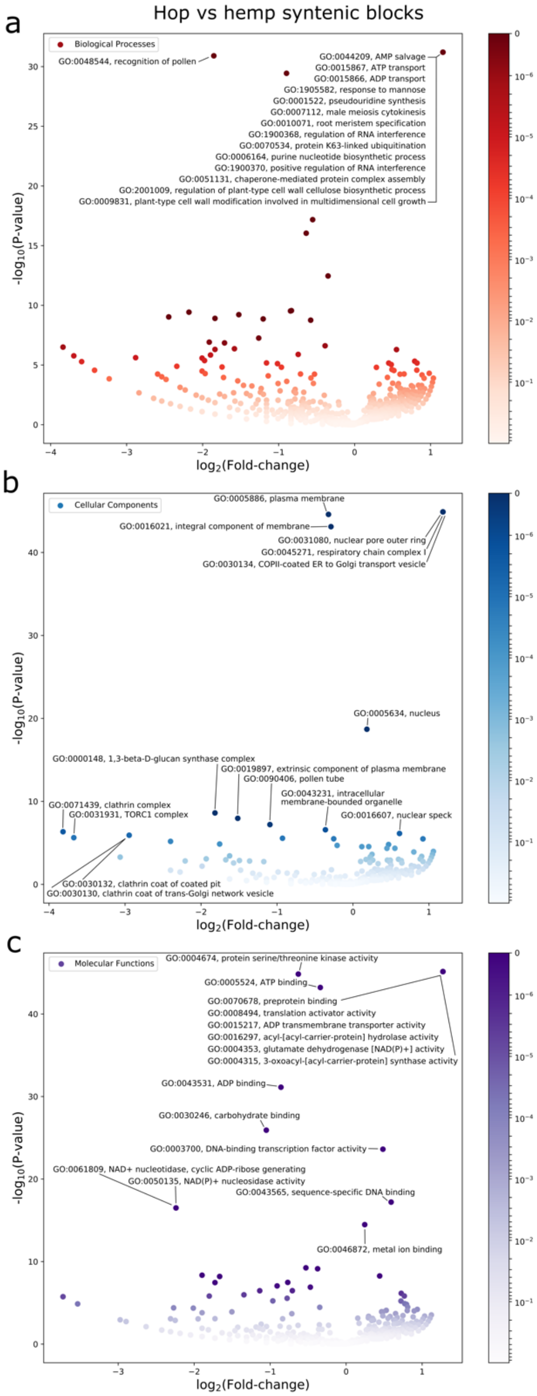

**Fig. S10. OrthoFinder results.** Bar charts showing the resulting statistics from OrthoFinder. **a)** Transdecoder gene models only; **b)** Transdecoder gene models and all Maker gene models; and **c)** Transdecoder gene models and Maker genes containing similarity to a known UniProt gene or Pfam domain. The three categories are shown for each of the eight species on the x-axis. The categories include the percentage of genes in all orthogroups, the percentage of orthogroups containing a given species, and the percentage of species-specific orthogroups. Our goal from this comparison was to find the set of gene models that maximized the number of orthogroups containing species, while minimizing species-specific orthogroups without similarity to a known gene.

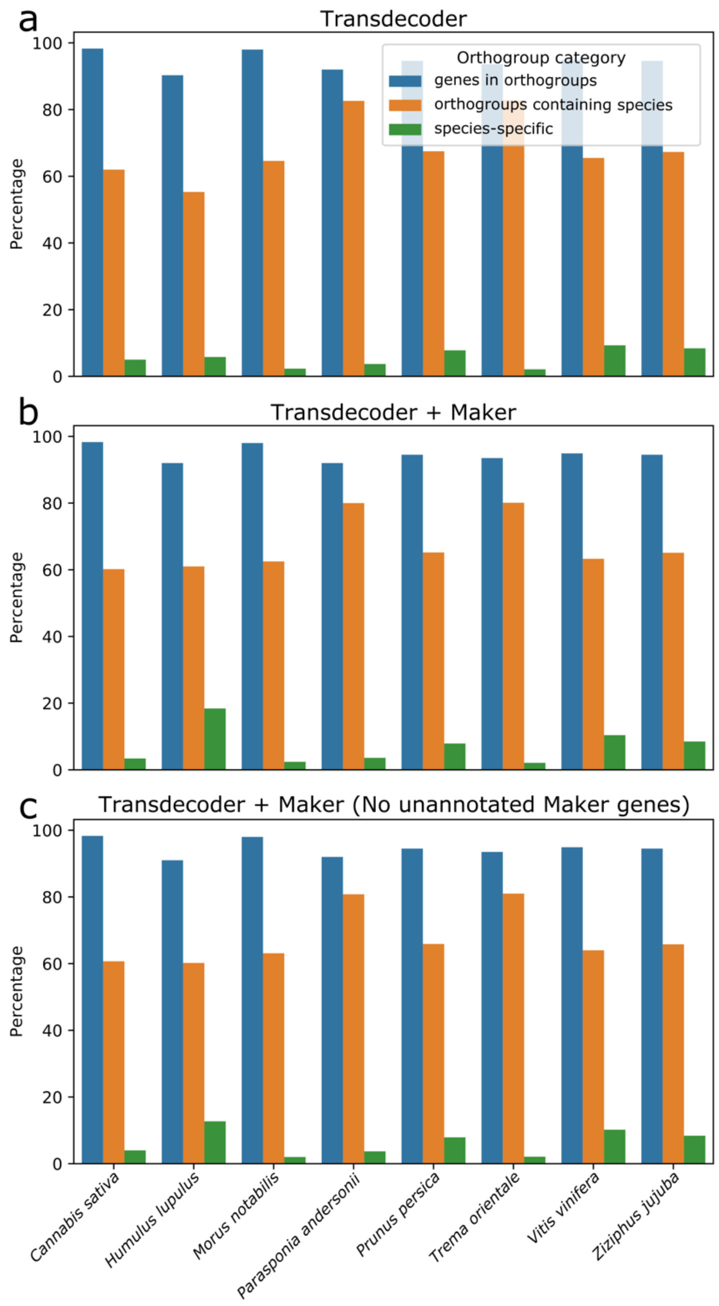

**Fig. S11. MCMCTree convergence. a)** Assessment of convergence by comparing the posterior means between two replicate runs for the strict molecular clock model (clock1). **b)** Assessment of convergence by comparing the posterior means between two replicate runs for the independent log-normally distributed relaxed-clock model (clock2). **c)** Trace of the posterior mean for the log-likelihood for the strict molecular clock. Translucent portion of trace denotes burn-in. The effective sample size (ESS) is 9,001. **d)** Trace of the posterior mean for the log-likelihood for the independent log-normally distributed relaxed-clock model. Translucent portion of trace denotes burn-in. The ESS is 8,885.

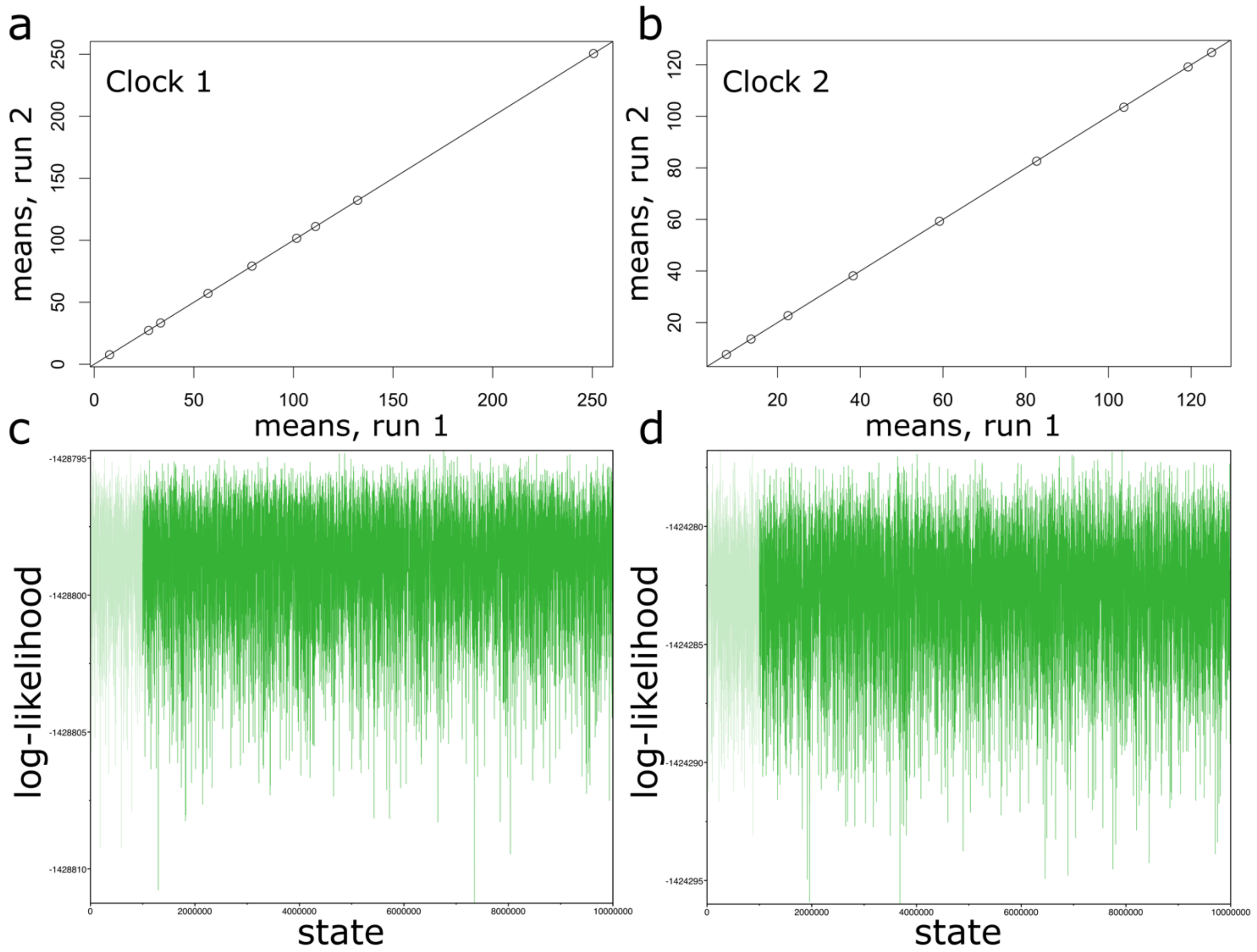

**Table S1.** Statistics about Hi-C libraries and HiRise assembly

| | |
|---|---|
| Number of breaks made to input assembly by HiRise | 1,027 |
| Number of joins made by HiRise | 8,131 |
| Library 1 stats | 156M read pairs; 2x150 bp |
| Library 2 stats | 92M read pairs; 2x150 bp |
| Library 3 stats | 143M read pairs; 2x150 bp |
| Library 4 stats | 67M read pairs; 2x150 bp |

**Table S2.** Analysis of polished assembly quality

| | |
|---|---|
| Substitution Errors | 961,874 bp |
| Insertion/Deletion Errors | 421,693 bp |
| Assembly Size | 3,712,781,139 bp |
| Consensus Quality | 99.9627% |

**Table S3.** Estimated genome sizes of *Humulus* and closely-related species

| Species or variety | Estimated genome size (pg) | Estimated haploid genome size (Gb) | Reference |
|---|---|---|---|
| *Humulus lupulus* var. *cordifolia* | 2.9 (1C) | 2.842 | (Zonneveld *et al.*, 2005; Leitch *et al.*, 2019) |
| *H. lupulus* var. *lupulus* cv. Lubelski | 5.598±0.044 (2C) | 2.743 | (Grabowska-Joachimiak *et al.*, 2006) |
| *H. lupulus* var. *lupulus* cv. Shinshu Wase | 5.62 (2C) | 2.57 | (Natsume *et al.*, 2015) |
| *H. lupulus* cv. Brambling Cross | 3.1 (1C) | 2.989 | (Zonneveld *et al.*, 2005) |
| *H. lupulus* var. *neomexicanus* | 6.064±0.048 (2C) | 2.971 | (Grabowska-Joachimiak *et al.*, 2006) |
| *H. scandens* (synonymous with *H. japonicus*) | 3.208±0.028 (Grabowska-Joachimiak *et al.*, 2006) (2C); 1.7 (Zonneveld *et al.*, 2005; Leitch *et al.*, 2019) (1C) | 1.572-1.666 | (Zonneveld *et al.*, 2005; Grabowska-Joachimiak *et al.*, 2006; Leitch *et al.*, 2019) |

**Table S4.** Hop genome heterozygosity and repeat content based on short-read DNA sequencing

| Results from GenomeScope 2.0 (ploidy=2; k-mer=21) | | |
|---|---|---|
| Property | Min | Max |
| Homozygous (aa) | 94.5323% | 95.4073% |
| Heterozygous (ab) | 4.59269% | 5.4677% |
| Genome Haploid Length | 1,572,209,199 bp | 1,658,246,355 bp |
| Genome Repeat Length | 1,207,266,703 bp | 1,273,332,843 |
| Genome Unique Length | 364,942,496 bp | 384,913,512 |
| Model Fit | 33.166% | 92.0062% |
| Read Error Rate | 0.480394% | |

**Table S5.** Linkage group statistics for the mapping population USDA 2017014

| Linkage Group | #Markers | Map Size | Average Gap Size | Biggest Gap Size | #Unique Positions |
|---|---|---|---|---|---|
|  |  | --------------------cM--------------------- |  |  |  |
| 1 | 487 | 128.419 | 0.26 | 2.731 | 487 |
| 2 | 311 | 94.254 | 0.3 | 2.505 | 311 |
| 3 | 327 | 125.717 | 0.39 | 3.786 | 326 |
| 4 | 406 | 121.541 | 0.3 | 1.435 | 405 |
| 5 | 520 | 110.051 | 0.21 | 0.838 | 520 |
| 6 | 365 | 119.981 | 0.33 | 2.983 | 365 |
| 7 | 209 | 164.823 | 0.79 | 15.049 | 209 |
| 8 | 627 | 84.314 | 0.13 | 0.421 | 627 |
| 9 | 399 | 135.052 | 0.34 | 1.735 | 398 |
| 10 | 439 | 185.377 | 0.42 | 1.513 | 438 |
| All | 4,090 | 1,269.529 | 0.35 | 15.049 | 4086 |

**Table S6. Transdecoder gene model results.**

| Transdecoder gene models | |
|---|---|
| **Category** | **Count** |
| Total gene models (complete + fragmented) | 57,684 |
| Complete protein-coding transcripts | 38,711 |
| Longest complete open reading frame (ORF) in full assembly | 21,698 |
| Longest complete open reading frame (ORF) in ten largest scaffolds | 20,581 |
| Lacking start codon | 11,691 |
| Lacking stop codon | 4,395 |
| Lacking start and stop codons | 2,887 |

**Table S7.** Gene model BUSCO results

| Category | Full assembly or largest 10 scaffolds | Database | % Total complete | % Single-copy complete | % Duplicated complete | % Fragmented | % Missing |
|---|---|---|---|---|---|---|---|
| Gene models (Transdecoder + Maker) | full | Embryophyta (1614) | 78.3 | 57.7 | 20.6 | 6.9 | 14.8 |
| **Gene models (Transdecoder + Maker)** | **10** | **Embryophyta (1614)** | **76.5** | **58.9** | **17.6** | **6.8** | **16.7** |
| Gene models (Transdecoder) | full | Embryophyta (1614) | 74.2 | 55.0 | 19.2 | 6.4 | 19.4 |
| Gene models (Transdecoder) | 10 | Embryophyta (1614) | 72.9 | 56.4 | 16.5 | 6.4 | 20.7 |

**Table S8. Number of genes with GO terms**

| **GO term category** | **Number of genes associated with GO term category** |
|---|---|
| Biological Processes | 19,147 |
| Cellular Components | 20,674 |
| Molecular Function | 19,385 |

**Table S9.** Pfam repeat-associated domains (this table is provided as a separate Excel file)

**Table S10.** Conditional repeat Pfam domains

| Pfam Accession ID | Pfam Accession Description |
|---|---|
| PF04434.18 | SWIM zinc finger |
| PF16588.6 | C2H2 zinc-finger |
| PF02892.16 | BED zinc finger |
| PF13917.7 | Zinc knuckle |
| PF14372.7 | Domain of unknown function (DUF4413) |
| PF09331.12 | Domain of unknown function (DUF1985) |
| PF13962.7 | Domain of unknown function |
| PF00385.25 | Chromo (CHRromatin Organisation MOdifier) domain |
| PF02902.20 | Ulp1 protease family, C-terminal catalytic domain |
| PF18907.1 | Family of unknown function (DUF5662) |
| PF00574.24 | Clp protease |
| PF11604.9 | Copper binding periplasmic protein CusF |
| PF13634.7 | Nucleoporin FG repeat region |
| PF00230.21 | Major intrinsic protein |
| PF05193.22 | Peptidase M16 inactive domain |

**Methods S1.**

**SNP identification and filtration**

To identify SNPs, we used the TASSEL 5.0 Pipeline (Glaubitz *et al.*, 2014) on default settings. The polished Cascade assembly was used as a reference genome for alignment with bowtie2 (Langmead & Salzberg, 2012). Approximately 1.3 million SNPs were identified and pre-filtered using default settings. The initial set of markers used for genetic mapping were filtered iteratively in TASSEL 5.52 (Bradbury *et al.*, 2007), alternating between incremental increases in stringency of genotypes or markers. Ultimately, a set of 55,214 markers across all samples (281 offspring and the two parental genotypes) were selected with 15% missing markers allowed, minor allele frequency of at least 0.2, and all selected genotypes having at least 80% presence across all samples. Missing marker data was then imputed using LD KNNi imputation (Money *et al.*, 2015).

The resulting data were then separated into separate pseudo-chromosomal groups and each group was exported as a VCF file.

**Haplotype-tagging SNPs (htSNPs) and SNPtag data**

SNPs from all 10 pseudo-chromosomes were split into individual sets with each set representing a single pseudo-chromosome. Pseudo-chromosome-specific data sets (in VCF format) were imported into JMP Genomics (JMP®, SAS Institute Inc. , Cary, NC, 1989–2021., 1989-2021) and converted to numeric data. We then identified linkage disequilibrium blocks (using “LD Block Creation”) for initial bin-formation. Maximum range of LD Block creations was set to “20 markers,” the Haplotype Estimation Model was set to “EST,” and the Block options were set as follows: MAF = 0.1; all other values are default. We subsequently ran, “Haplotype Estimation” on the resulting data sets from LD Block Creation and used the resulting data sets for identification of haplotype-tagging SNPs (“htSNP analysis”). The two output files resulting from “htSNP” analyses consisted of one file containing all possible SNPs—both those representing bins as well as SNPs not located within a linkage block (singletons)—and a file with just SNPs representing LD bins. The latter file was then filtered to have just one representative SNP per LD bin (SNPtags). This filtered file was subsequently used for development of the genetic maps for each of the pseudo-chromosomes with the exception of pseudo-chromosome 7. In this one case, the resulting data set exhibited fewer linkage blocks that were larger on average than the other pseudo-chromosomes. As a result, we added singletons to the starting file for development of a genetic map for this linkage group.

**Genome size and heterozygosity**

We used KmerGenie version 1.7051 to perform the k-mer distribution analysis, which determined an optimal k-mer size of 75 (Chikhi & Medvedev, 2014). We also estimated the heterozygosity of the genome using GenomeScope 2.0, which recommends a k-mer size of 21 for most genomes (Ranallo-Benavidez *et al.*, 2020). We used KMC (Kokot *et al.*, 2017) to count k-mers as input for GenomeScope 2.0. To perform these analyses, we used the short-read DNA sequencing from Cascade.

**RNA extraction and processing for RNA-seq**

RNA was extracted from gland and leaf hop tissues using a Qiagen RNeasy kit. The lysis buffer in the kit was replaced with a buffer of 4M guanidine isothiocyanate, 0.2 M sodium acetate pH 5.0, 25mM EDTA, 2.5% (w/v) PVP-40, and 1% (v/v) Beta-mercaptoethanol. The quality of the RNA was tested on an Agilent Bio-analyzer. The extracted total RNA passing QC was used in an Illumina TruSeq RNA library preparation kit. The resulting cDNA libraries were sequenced on an Illumina HiSeq at Illumina's California campus.

**Gene model development with Transdecoder**

To predict protein-coding transcripts, we used Transdecoder-v5.5.0 (Furuno *et al.*, 2003; Haas *et al.*, 2013). We aligned RNA-seq from lupulin glands, leaf, meristem, stem tissues, as well as hop cones during critical developmental stages (Padgitt-Cobb *et al.*, 2021; Eriksen *et al.*, 2021) to the Cascade Dovetail assembly, and then assembled transcripts. Alignment of RNA-seq to the assembly was performed with hisat2 version 2.2.0 (Kim *et al.*, 2019; Xu *et al.*, 2020), followed by transcript assembly with StringTie v1.3.3b (Pertea *et al.*, 2015; Xu *et al.*, 2020), and finally, merging of transcript-specific assemblies with the 'stringtie --merge' option.

Transdecoder identifies the coding DNA sequence (CDS) for each transcript from the longest open reading frame (ORF) (Furuno *et al.*, 2003; Xu *et al.*, 2010). Transdecoder makes its decision by incorporating the length of the open reading frame (ORF), a log-likelihood score, and a position-specific scoring matrix (PSSM) to refine the transcript boundaries. Transdecoder categorizes ORFs into four types, depending on the presence of start and stop codons, including "complete," "5prime_partial," "3prime_partial," and "internal." Transcripts identified as "5prime_partial" do not have a start codon, "3prime_partial" do not have a stop codon, and 'internal' do not contain start or stop codons.

We extracted transcripts from the genome assembly with the following command: "TransDecoder-v5.5.0/util/gtf_genome_to_cdna_fasta.pl stringtie_merged.gtf cascadeAssembly.fasta > transcripts.fasta." We generated a corresponding GFF file with the following command: “TransDecoder-v5.5.0/util/gtf_to_alignment_gff3.pl stringtie_merged.gtf > transcripts.gff3.”

Transdecoder proceeds in multiple steps. First, we extracted the longest ORF sequences by running the command, “TransDecoder.LongOrfs -S -t transcripts.fasta.” We then identified transcripts with similarity to UniProt Embryophyta protein sequences (38,747 sequences, accessed 08/24/2020) and Pfam domains (Pfam release 33.1). Next, we predicted genes with the command, “TransDecoder.Predict -t transcripts.fasta --retain_pfam_hits pfam.domtblout --retain_blastp_hits uniprot.blastp.” We generated a genome coordinate-centric GFF file with the following command:
“TransDecoder-v5.5.0/util/cdna_alignment_orf_to_genome_orf.pl transdecoder.gff3 transcripts.gff3 transcripts.fasta > transdecoder.genomeCentric.gff3.” For all subsequent analyses, we used the longest ORF per transcript. A minimum protein length of 100 amino acids was required, which is also the default length.

**Gene model development with MAKER**

The first step toward developing the set of gene models with MAKER involved generating alignment evidence. We used megablast 2.2.26 (Altschul *et al.*, 1990) to align ESTs from NCBI (25,692 sequences, accessed 11/12/2018) and TrichOME (Dai *et al.*, 2010) (22,959 sequences, accessed 03/28/2018) to the assembly. After obtaining initial alignments with megablast, we collected ESTs that aligned to the assembly, and performed a secondary alignment with est2genome (exonerate version 2.3.0) (Slater & Birney, 2005). We also generated protein alignments with *Cannabis sativa* (RefSeq; 33,639 sequences, accessed accessed 12/02/2020), *Prunus persica* (PLAZA (Van Bel *et al.*, 2018); 26,843 sequences, accessed 10/13/2020), *Ziziphus jujuba* (PLAZA (Van Bel *et al.*, 2018); 28,799 sequences, accessed 10/13/2020), and Embryophyta protein sequences (UniProt; 38,747 sequences, accessed 08/24/2020). For the protein sequences, we performed a first round of alignment with blastx 2.10.0+ and then performed a second round of alignment with exonerate protein2genome (exonerate version 2.3.0) (Slater & Birney, 2005) on the sequences that aligned to the assembly with blastx.

We also generated transcripts from RNA-seq for gene model prediction. Our approach began with alignment of RNA-seq to the assembly with hisat2 version 2.2.0 (Kim *et al.*, 2019; Xu *et al.*, 2020), followed by transcript assembly with StringTie v1.3.3b (Pertea *et al.*, 2015; Xu *et al.*, 2020), and finally, merging of transcript-specific assemblies from leaf, meristem, and stem tissues with cuffmerge 2011-03-17 (Trapnell *et al.*, 2010).

We generated final consensus gene models with MAKER (Cantarel *et al.*, 2008; Holt & Yandell, 2011; Campbell *et al.*, 2014; Xia *et al.*, 2019), using both Augustus-3.3.2 (Stanke *et al.*, 2006) and SNAP (Korf, 2004; Xu *et al.*, 2020) to perform gene model prediction. We trained Augustus by running BUSCO v4.1.1 in 'long' mode (Waterhouse *et al.*, 2018; Elbers *et al.*, 2019; Bohn *et al.*, 2021). Gene model development with MAKER proceeded in three steps. For the first round of MAKER, only the alignments were included as evidence in the config file (est2genome=1; protein2genome=1), to generate an initial set of models for training with SNAP. After the first round of MAKER, we trained SNAP. The resulting "snaphmm" file from SNAP was included in the second round of MAKER (est2genome=0; protein2genome=0). After the second round of MAKER, we trained SNAP again, producing an updated ''snaphmm" file. The third round of MAKER included both the second-round "snaphmm" file from SNAP, as well as the Augustus models trained by BUSCO (De-la-Cruz *et al.*, 2021).

We used gffread to extract CDS and full transcripts (Pertea & Pertea, 2020). The detailed pipeline for gene prediction, including commands and MAKER config files, can be found at GeneModels/MakerGeneDevelopmentPipeline.md on the GitHub project page (https://github.com/padgittl/CascadeDovetail).

**Density of genes and long terminal retrotransposons (LTRs)**

We calculated the density of genes and LTRs by counting the occurrence of gene and LTR sequences in a 5 Mb window.

**Gene model homology**

For evolutionary and orthology analyses, we obtained gene models from NCBI for *Cannabis sativa* (RefSeq assembly accession GCF_900626175.2), *Morus notabilis* (RefSeq assembly accession

GCF_000414095.1), *Parasponia andersonii* (GenBank assembly accession GCA_002914805.1), *Trema orientale* (GenBank assembly accession GCA_002914845.1), and *Vitis vinifera* (RefSeq assembly accession GCF_000003745.3). We first removed genes annotated as pseudogenes or low quality and then included only the longest transcript per gene. We obtained the "selected transcripts" for *Prunus persica* and *Ziziphus jujuba* from PLAZA (accessed 10/13/2020), corresponding to the longest transcript.

For defense and disease response-associated genes, we collected GO terms (Ashburner *et al.*, 2000) including keywords 'defense' or 'disease,' and then retrieved all UniProt genes associated with the selected GO terms.

**Repeat-associated gene models**

We followed an adapted version of our previously described pipeline to identify repeat-associated genes using both Pfam domains (Finn *et al.*, 2014) and UniProt genes. First, we collected 164 transposable element (TE)- and virus-associated Pfam domains (Pfam release 33.1) based on keyword search and literature (Supplementary Table 9). From UniProt, we downloaded sets of genes from bacteria (accessed 04/15/2021 using search term: taxonomy:"Bacteria [2]" AND reviewed:yes), viruses (accessed 04/15/2021 using search term: taxonomy:"Viruses [10239]" AND reviewed:yes), and TEs (accessed 04/15/2021 using search term: keyword:"Transposable element [KW-0814]" AND reviewed:yes). We also collected 12 transposon-associated genes from the set of 38,747 UniProt Embryophyta genes (accessed 08/24/2020 using search term: taxonomy:"Embryophyta [3193]" AND reviewed:yes).

We aligned the gene models to each set of UniProt genes in both directions using blastp (version 2.12.0+), applying an E-value threshold of less than 1e-3. We aligned the gene models to the Pfam domains with hmmscan (HMMER 3.3 (Nov 2019)), applying an E-value threshold of less than 1e-3.

We identified three categories of genes with similarity to Pfam domains: similarity only to non-repeat-associated domains, similarity only to repeat-associated domains, and similarity to both repeat- and non-repeat-associated domains. We further split the set of genes with similarity to both repeat- and non-repeat-associated domains into two categories. For the first category, we identified genes with domains that frequently occurred with repeat domains, but did not have apparent repeat similarity when they occurred alone. We designated this set of domains as "conditional repeats" (Supplementary Table 10). For the remaining examples with both repeat- and non-repeat Pfam domains, without "conditional repeat" domains, we applied a coverage filter. If more than 30% of the length of a gene was similar to repeat domains, that gene was reassigned as a repeat gene (Ye & Zhong, 2015; Yang *et al.*, 2016).

To assign similarity to UniProt genes, we required a minimum percent identity based on alignment length, known as the "Twilight zone" of protein alignment (Rost, 1999). For Embryophyta UniProt genes, we also imposed a minimum query coverage of 20%, and for bacteria, virus, and TE UniProt genes, we imposed a minimum query coverage of 30%.

The final step in the filtering pipeline involved compiling all of the sources of alignment evidence to assign genes to one of three categories: non-repeat, repeat, or unannotated. First, to assign a gene as non-repeat, we checked if genes with similarity to UniProt Embryophyta genes did not also have only similarity to Pfam repeat domains. If all domain similarity was repeat-associated, the gene was assigned to the repeat category.

For genes with similarity to a UniProt TE gene, we checked first if these genes had similarity to a UniProt Embryophyta gene. If so, we required the gene with similarity to a TE to have greater query coverage to the TE than to the Embryophyta gene, to be reassigned as repeat.

For genes with similarity to UniProt bacteria and virus genes, we required that the gene did not already have similarity to an Embryophyta gene, to avoid removing genes with ancient conserved function. Finally, we collected genes lacking any similarity to a UniProt gene or Pfam domain. We applied this pipeline to the set of hop gene models, as well as the other seven species in our orthology analysis. The detailed pipeline for gene annotation can be found at GeneModels/GeneAnnotationPipeline.md on the GitHub project page (https://github.com/padgittl/CascadeDovetail).

**Calculation of 4DTv**

The rate of transversions at the third position in four-fold degenerate (4D) codons (including amino acids A, G, L, P, R, S, T, V) (Tang *et al.*, 2008) can be used as a simple molecular clock, and to assess the occurrence of large-scale duplication events (Hellsten *et al.*, 2007; Xiao *et al.*, 2015). Transversions occur at a slower rate than transitions; the accumulation of synonymous substitutions at the third position in 4D codons provides a measure of neutral genetic divergence (Hellsten *et al.*, 2007; Rozenfeld *et al.*, 2019). The 4DTv rate is less susceptible to saturation effects (Gojobori, 1983; Smith & Smith, 1996) than Ks (Potato Genome Sequencing Consortium *et al.*, 2011; Unver *et al.*, 2017; Schiavinato *et al.*, 2020; Shingate *et al.*, 2020). The range of 4DTv is from 0.0, corresponding to recent duplication, to approximately 0.5, where sequences are sufficiently diverged to be practically random (Hellsten *et al.*, 2007). We required uncorrected 4DTv values to be less than 0.5 (Rozenfeld *et al.*, 2019).

We performed a codon-level alignment for each anchor gene pair using MACSE alignSequences and exportAlignment (Ranwez *et al.*, 2018), and calculated 4DTv distances for each collinear gene pair individually (Xiao *et al.*, 2015), similar to the procedure for calculating Ks. To calculate 4DTv distances, we used a custom Python script (see MolecularEvolution/scripts/calculate4DTv.py on our GitHub project page). For the 4DTv calculation, we required a minimum of 25 4D codons shared between sequences in the alignment, and corrected for multiple substitutions (Hellsten *et al.*, 2007; Tang *et al.*, 2008).

**Fossil calibration dates**

We used a fossil calibration date of 5-28 mya based on previously described calibration intervals from literature (Tiffney, 1986; Collinson, 1989; Zerega *et al.*, 2005; McPartland, 2018; Jin *et al.*, 2020). Most recently, Jin *et al.* restricted the *Humulus* constrained to 23 mya based on fruit fossil evidence of *Humulago reticulata* (Dorofeev) Doweld from Antropovo, Russia during the Oligocene (Collinson, 1989; Doweld, 2016), resulting in an estimated divergence date of 25.4 mya for *Humulus* and *Cannabis* (Jin *et al.*, 2020). Zerega *et al.* also set the fossil calibration date to 5-23 mya for *Humulus*, placing the divergence of *Humulus* and *Cannabis* at 21 mya (Zerega *et al.*, 2005). McPartland (McPartland, 2018) used a calibration date of 16-28 mya, placing *Humulus* and *Cannabis* divergence at the boundary of the early and late Oligocene (28 mya) based on morphological evidence for a "generalized" dispersal mechanism in *Humulus* and *Cannabis*

(Tiffney, 1986). Calibration dates for *Trema* and *Parasponia* based on fossil evidence are not available, to our knowledge.

We applied calibration dates for species outside of the Cannabaceae from TimeTree.org (Kumar *et al.*, 2017), while also verifying these dates with literature. He *et al.* estimated the divergence date of 63.5 mya for mulberry (*Morus notabilis*) and *Cannabis sativa* (He *et al.*, 2013). Early fossil evidence for the *Moraceae* family dates from the Eocene (56-33.9 mya). Based on molecular dating, the Rosidae clade has an estimated origination date between 123–93 mya (Magallón *et al.*, 2015; Sun *et al.*, 2016), and the estimated date for the divergence of *Vitis* is 115 mya (Fawcett *et al.*, 2009). The emergence of the eudicots is estimated to have occurred around 125 mya, and is considered to be a reliable calibration date for molecular dating analyses (Doyle *et al.*, 1977; Hickey & Doyle, 1977; Doyle & Hotton, 1991; Fawcett *et al.*, 2009; Xiang *et al.*, 2017).

**MCMCTree parameters**

We used a likelihood-ratio test (LRT) to compare molecular clock models (Yang *et al.*, 2000) with the equation LRT=-2(ln(Ls)-ln(Lg)). L is the likelihood value, ln(x) is the natural logarithm, and s corresponds to the simpler model with fewer parameters than the general model g. We used the HKY85 substitution model (model=4) (Hasegawa *et al.*, 1985) and set the root age to <1.25, corresponding to 125 mya. Other parameters for MCMCTree include burnin = 50,000, sampfreq = 1,000, and nsample = 10,000. We evaluated the MCMC results with Tracer v1.7.2 (Fig. S11) (Suchard *et al.*, 2018; Dos Reis & Yang, 2019). The control file for MCMCTree can be viewed at the GitHub page under 'TimeDivergenceEstimation'.
